# Satellite Tobacco Mosaic Virus: Revealing Environmental Drivers of Capsid and Nucleocapsid Plasticity using High-Resolution Simulations

**DOI:** 10.1101/2025.01.28.635309

**Authors:** Marharyta Blazhynska, Olivier Adjoua, Zoé Vetter, Louis Lagardère, Jean-Philip Piquemal

## Abstract

The Satellite Tobacco Mosaic Virus (STMV) serves as a model system for elucidating how electrostatic and mechanical forces shape single-stranded (ss) RNA viral architecture. Leveraging a cumulative total of 1.5 *µ*s of simulation with a polarizable force field including 1.2 *µ*s of conventional Molecular Dynamics (MD) supplemented by Gaussian accelerated MD (GaMD), and well-tempered metadynamics (WTMetaD) enhanced sampling techniques, we examined how pH and ionic composition regulate the structural dynamics of preassembled STMV capsids. Six ∼1M-atom assemblies spanning physiological and stress-mimicking environments were modeled to capture the interplay among protein–protein, protein–RNA, and ion-mediated interactions. A representative GaMD trajectory, corresponding to the up to *µ*s-equivalent regime of conventional MD, revealed that the capsid undergoes coordinated radial fluctuations, with collective expansion and contraction of the icosahedral shell. WTMetaD free-energy surfaces, computed for all six assemblies, delineated distinct condition-specific minima, defining thermodynamically accessible conformations for each environment. During the early relaxation dynamics revealed in conventional MD, RNA-free capsids preserved their icosahedral symmetry in physiological salt, where monovalent ions screened intra-capsid electrostatics. RNA encapsidation further modulated this balance through divalent-ion coordination at the protein–RNA interface. Under acidic conditions, a reversible Na^+^/Mg^2+^ exchange reorganized interfacial charge networks while maintaining the overall capsid architecture. Transient chloride binding intermittently disrupted key inter-monomer salt bridges, exposing a regulatory mechanism for capsid plasticity and permeability. Our findings establish STMV as an inherently dynamic, ion-responsive structural assembly whose conformational adaptability emerges from finely balanced electrostatic coupling between the capsid and its RNA, providing an atomistic framework for how ssRNA icosahedral viruses sense and adjust to environmental changes.

## Introduction

The Satellite Tobacco Mosaic Virus (STMV) is a single-stranded RNA (ssRNA) virus with a T=1 icosahedral capsid of approximately 170 Å in diameter^1,2^. STMV relies on tobacco mosaic virus as a replication helper and serves as a benchmark system for dissecting the molecular principles underlying viral assembly^3–6^. While its architecture has been resolved to atomic precision, the molecular mechanisms governing its dynamic stability, particularly how the capsid and RNA respond to physicochemical perturbations, remain poorly understood^1,2,7–10^.

Viral capsids are stabilized by a delicate balance of inter-monomer forces, including hydrophobic contacts, hydrogen bonding, and electrostatic networks, which collectively define the geometric organization of the icosahedral shell^11–13^. In many ssRNA viruses, encapsidated RNA modulates the capsid geometry through charge compensation and salt-bridge formation with protein residues. Experimental studies demonstrated that STMV and related ssRNA viruses can form RNA-free capsids that preserve near-native icosahedral symmetry under suitable ionic and pH conditions^5,11–19^. In contrast, classical all-atom molecular dynamics (MD) simulations have historically struggled to reproduce this behavior. Pioneering studies by Schulten et al.^20,21^ showed that empty STMV capsids tend to collapse within a few nanoseconds, whereas RNA-filled capsids remained intact. These discrepancies between computational and experimental studies motivated a systematic reassessment of STMV-related assemblies. Although more recent high-resolution structures exist^22,23^, we deliberately reproduced the original Schulten et al. models to enable a one-to-one comparison between classical and polarizable force fields under identical starting conditions.

Building on this baseline, we extended our investigation by introducing six biologically relevant scenarios to capture the range of physicochemical environments that STMV may potentially encounter during its life cycle^24^. Our goal was to systematically probe how variations in pH, ionic strength, and cation composition influence capsid architecture, RNA–protein electrostatics, and global conformational adaptability under physiologically relevant and stress-mimicking conditions^25–35^. The component choice is further supported by *in vivo* and *in situ* studies on plant vacuolar ion accumulation and pH regulation^36–39^. The investigated assemblies were as follows:

1. Empty STMV capsid neutralized with chloride ions and pH=7, reproducing Schulten et al. original study.
2. Empty STMV capsid at physiological NaCl concentration (i.e., 0.15 M)^40^ and pH=7, representing cytoplasmic-like conditions to evaluate the intrinsic conformational behavior of the empty shell.
3. STMV nucleocapsid at physiological concentration, providing a baseline for the influence of the RNA genome on capsid geometry and interfacial interactions in the absence of divalent cations.
4. STMV capsid neutralized with Mg^2+^ and Cl^−^ ions, reproducing Schulten et al. nucleocapsid scenario.
5. STMV nucleocapsid at 0.15 M NaCl supplemented with Mg^2+^ at pH=7, mimicking cytoplasmic ionic conditions where magnesium ions are expected to reinforce RNA–protein electrostatic interactions.
6. RNA-bound nucleocapsid at 0.5 M NaCl with Mg^2+^ at pH=5, capturing acidic and highionic-strength intracellular compartments, such as vacuoles and vesicular trafficking pathways, relevant for viral life cycle and host interactions^29,30,33,34,41,42^.

The details of the studied STMV assembly conditions, including the number of water molecules, ions, and total atom count, are provided in Table 1. To our knowledge, these simulations represent the first atomistic exploration of how environmental conditions govern the structure and dynamics of STMV assemblies using fully polarizable MD. In this context, polarizable force fields provide a more detailed and accurate representation of molecular dynamics^43–45^, whereas classical all-atom non-polarizable MD approaches may fail to capture critical mechanisms of charge redistribution and subtle many-body intermolecular interactions^46^, leaving certain aspects of the behavior of large STMV-like assemblies underexplored. In particular, a polarizable force field such as AMOEBA^47–49^ is fully flexible and the combination of fully polarizable/flexible protein and water^50^ models permit the adjustment of dipole moments within the dynamic and have been shown to be of critical importance to match experiments when studying viral components, for example in the context of the SARS-CoV2 virus.^51–54^ In total, 1.5 *µ*s of fully polarizable MD trajectories were generated, substantially exceeding the sampling achieved in previous classical all-atom studies^20,21^. Within the conventional 100-ns trajectories, the capsid exhibited spontaneous, slow collective rearrangements, including radial expansion and contraction, providing mechanistic insight into the early pathways of capsid relaxation^55,56^. These motions, arising from cooperative protein–protein and RNA–protein interactions, were reproducible across independent triplicate trajectories, revealing emergent behaviors inherent to the intact viral assembly. Complementary dual-boost GaMD simulation (i.e., both potential and dihedrals of the whole system), equivalent of up to a *µ*s conventional MD regime^57–60^, on the empty STMV capsid under physiological conditions (№ 2, Table 1) reproduced the collective motions observed in the conventional MD. During GaMD, the capsid retained its overall icosahedral architecture while exhibiting continuous large-scale radial fluctuations, reflecting the intrinsic dynamic adaptability of icosahedral viral particles^61,62^. To quantify conformational behavior across the studied environments, quadruplicate well-tempered metady-namics (WTMetaD)^63–65^ simulations were performed, providing a thermodynamic description of the distinct conformational states sampled by each STMV assembly. The resulting free-energy surfaces converged to well-defined potential of mean force (PMF) minima, providing a robust thermodynamic description of the distinct conformational states sampled by each STMV assembly, highlighting the impact of environmental conditions on capsid dynamics. Collectively, our study provides a detailed, atomistic-level characterization of STMV capsid behavior, RNA–protein interactions, and ion-mediated modulation across a range of physiologically relevant environments. By systematically varying pH, ionic strength, and ion composition, and combining fully polarizable conventional MD with enhanced sampling approaches (GaMD and WTMetaD), we reveal how viral assemblies access distinct conformational states, adjust interfacial interactions, and couple RNA–protein motions in response to environmental cues, providing a mechanistic framework for understanding its environmental adaptability.

**Table 1.** Summary of Studied STMV Assemblies and System Compositions.

| N<sup>o</sup> | STMV Assembly | $N_{\text{atoms}}^{\text{tot}}$ | $N_{\text{water}}$ | $N_{\text{Mg}^{2+}}$ | $N_{\text{Na}^+}$ | $N_{\text{Cl}^-}$ |
| --- | --- | --- | --- | --- | --- | --- |
| 1* | Capsid neutralized, pH=7 | 1034475 | 299405 | – | – | 299 |
| 2* | Capsid in C(NaCl)=0.15M, pH=7 | 457162 | 106766 | – | 302 | 601 |
| 3* | Nucleocapsid in C(NaCl)=0.15M, pH=7 | 790181 | 207413 | – | 1066 | 586 |
| 4 | Nucleocapsid neutralized $\text{MgCl}_2$, pH=7 | 1066622 | 299854 | 474 | – | 299 |
| 5 | Nucleocapsid C(NaCl)=0.15M, $\text{Mg}^{2+}$, pH=7 | 789604 | 206953 | 474 | 841 | 1140 |
| 6 | Nucleocapsid C(NaCl)=0.5M, $\text{Mg}^{2+}$, pH=5 | 913572 | 247268 | 474 | 2358 | 2586 |

\*Conventional 100-ns MD simulations were performed in triplicate; only analysis of the first replica is presented in the main text. Root mean square deviation (RMSD) and gyration radius profiles (with standard deviations) for all replicas are provided in the Supporting Information (SI).

## Results and Discussion

### Capsid Structure and Dynamics

The structural organization of the STMV capsid and its interactions with encapsidated RNA are illustrated in Fig. 1A, B. The capsid, composed of 60 identical monomers (Fig. 1D), forming its icosahedral symmetry. Such geometric arrangement inherently creates structural voids — interstitial spaces between monomers^29,66^(Fig. 1C). Within each monomer (Fig. 1D), charged residues (i.e., ARG, LYS, ASP, and GLU) play a key role in both inter-monomer interactions and RNA binding.

**Figure 1.**
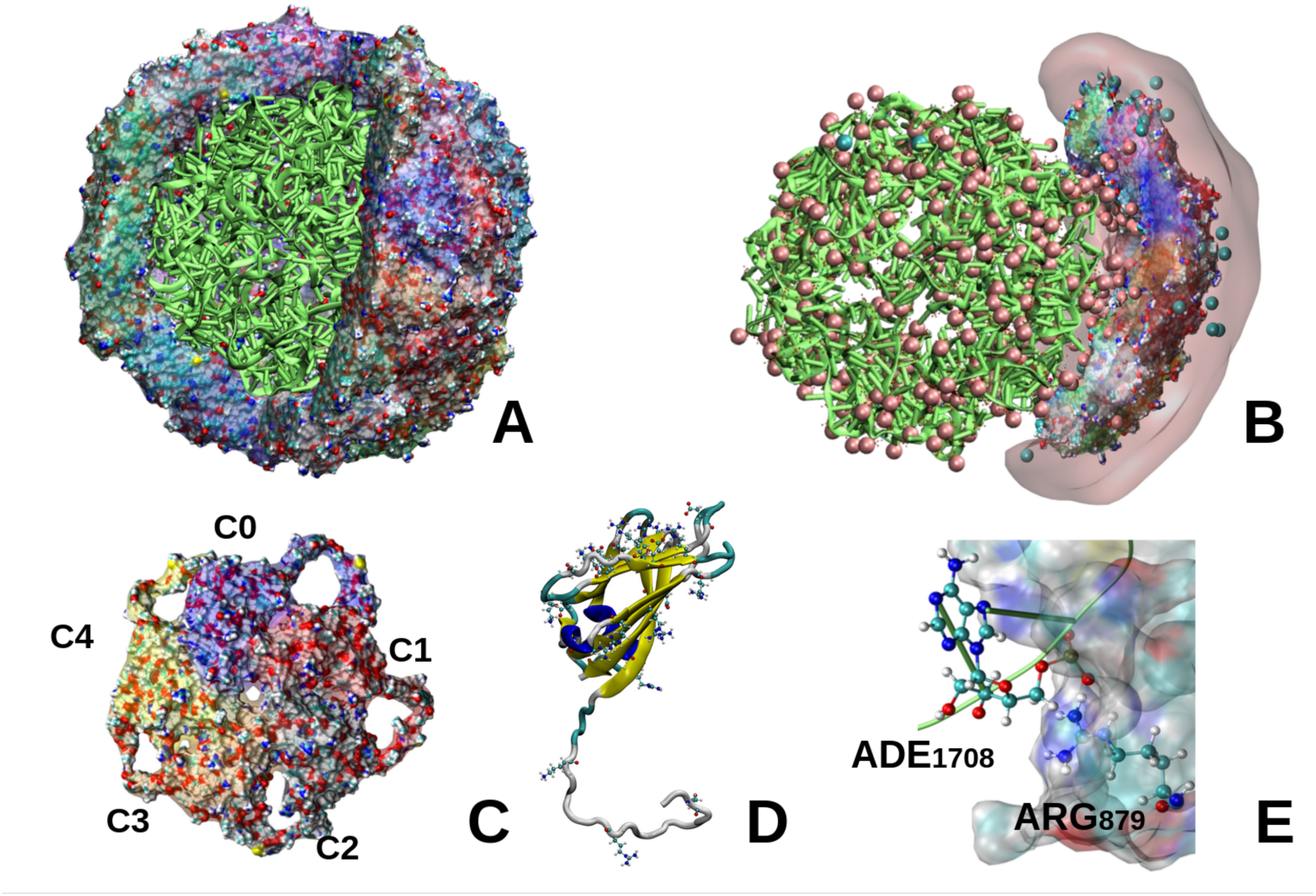
Structural features of STMV capsid and its interactions with RNA visualized employing Visual Molecular Dynamics software (VMD)^67^. (A) Molecular surface representation of the STMV capsid, colored by electrostatic potential^68^, with the encapsidated RNA shown shown in green. (B) Capsid–RNA complex with surrounding ions (Mg^2+^ and Cl^−^ in pink and teal spheres, respectively). (C) Surface representation of a pentameric capsid unit (C0–C4). (D) A representative capsid monomer unit with visualized charged residues of aspartic and glutamic acids, arginine, and lysine (ASP, GLU, ARG, and LYS). *β*-sheets, *α*-helices, loops, and turns are depicted in yellow, blue, white, and cyan, respectively. (E) Close-up of a RNA–protein interaction site, showcasing the most frequent interactions, occurring between the phosphodiester of RNA and protein sidechain on the example of ADE_1708_ and ARG_879_.

To probe the capsid structural dynamics, we analyzed the RMSD, radius of gyration (r_gyr_), root mean square fluctuation (RMSF) of C*α* atoms, as well as solvent-accessible surface area (SASA) of the capsid as a whole over 100–ns conventional MD simulations (Fig. 2A-D). To gain deeper insight into observed changes in capsid symmetry along the simulations, we additionally decomposed r_gyr_ along its principal axes, revealing directions of capsid anisotropic deformations (see details in Fig. S5 in SI). To identify monomeric regions involved in structural reorganization, we computed the mean RMSF of C*α* atoms per monomer along with its standard deviation (see Fig. S6), capturing both average flexibility and fluctuation variability across defined conditions (Table 1). Additionally, we examined the distribution of RMSF values across all C*α* atoms to assess whether fluctuations were broadly spread across the capsid or concentrated in specific regions. Time-dependent metrics continued to fluctuate throughout the 100-ns trajectories, reflecting the intrinsic non-stationary, breathing behavior of STMV rather than lack of equilibration, thus, capturing key aspects of the capsid dynamics and their sensitivity to ionic conditions.

**Figure 2.**
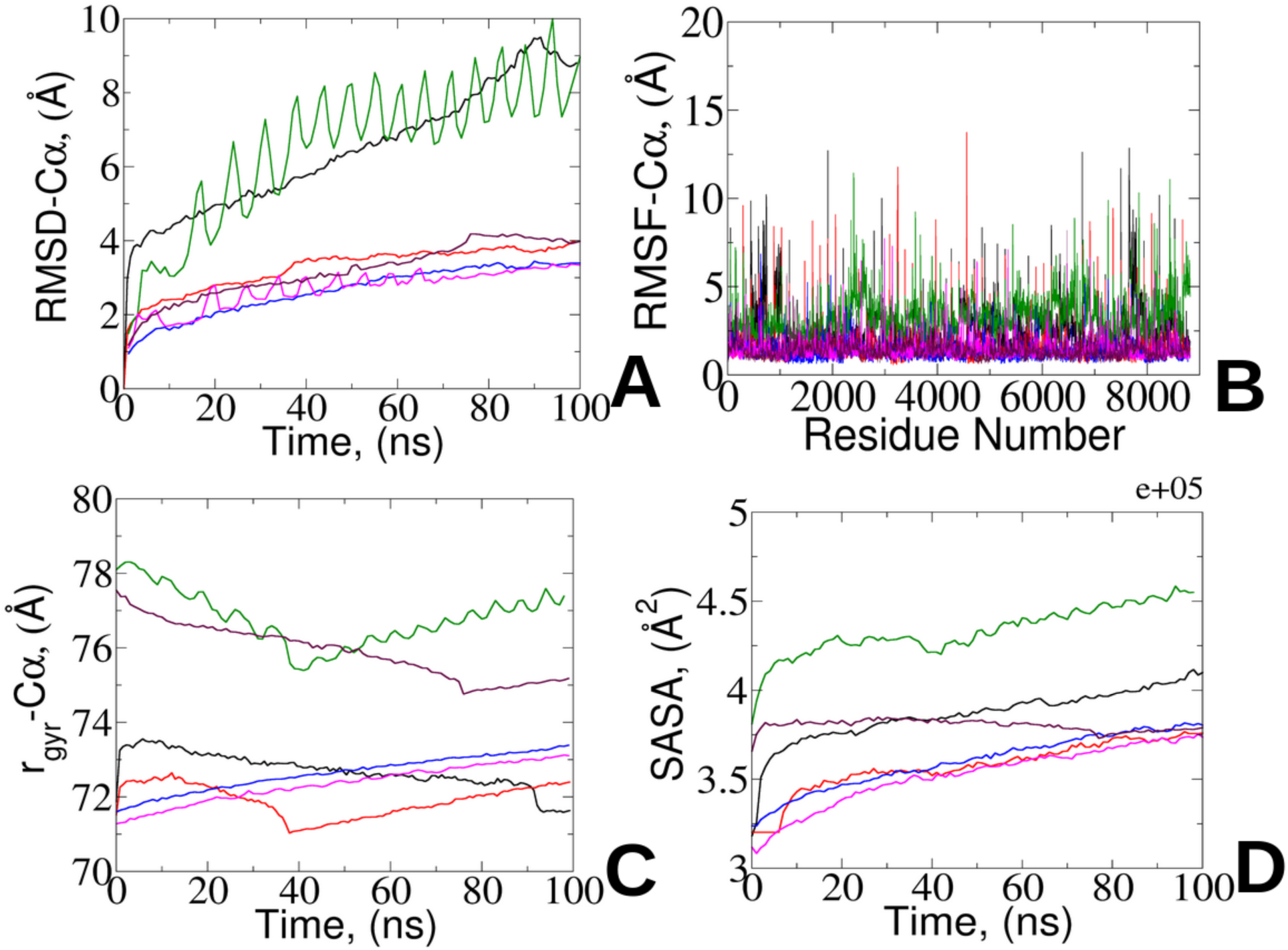
Analysis of the STMV capsid structural dynamics along conventional 100-ns-MD simulations: (A) Root Mean Square Deviation (RMSD) of C*α* atoms, (B) Root Mean Square Fluctuation (RMSF) of capsid C*α* atoms, (C) Capsid radius of gyration (r_gyr_), (D) Capsid solvent accessible surface area (SASA) across the different STMV conditions. The corresponding colors for each studied viral conditions are: № 1 - neutralized capsid with Cl^−^ ions (black), № 2 - capsid in physiological NaCl concentration (red), № 3 - nucleocapsid in physiological NaCl concentration (green), № 4 - nucleocapsid neutralized with MgCl_2_ (blue), № 5 - nucleocapsid in physiological NaCl concentration with Mg^2+^ present (magenta), and № 6 - nucleocapsid in 0.5 M NaCl concentration with Mg^2+^ mimicking pH=5 conditions (brown).

For the neutralized empty capsid, we observed one of the most pronounced changes in the global structure. Its RMSD reached ∼9 Å by the end of 100 ns, signifying substantial capsid reorganization (Fig. 2A, № 1). Although these rearrangements are more prominent than under other conditions, they do not lead to a rapid collapse, contrasting with earlier MD studies by Freddolino *et al*^20^. Instead, the capsid retains its overall architecture, albeit with noticeable reduction in global r_gyr_ over time (Fig. 2C). Decomposition of r_gyr_ along the principal axes reveals that this reduction is mostly pronounced along the *z*-axis, resulting in a slightly flatter capsid shape (Fig. S5). The RMSF analysis of C*α* atoms indicated that, while the most of fluctuation is in range of 2-6 Å (Fig. 2B), yet certain monomers (e.g., C3–C6 and C52) exhibit even higher fluctuations (extending up to 12 Å) with increased variability across these monomers (Fig. S6), suggesting zones of non-uniform rearrangements across the capsid. Additionally, the SASA steadily increases

(Fig. 2D), suggesting that the observed structural rearrangement might lead to a formation of additional interstitial voids. Remarkably, the inclusion of physiological NaCl (№ 2) moderated these changes (RMSD ∼3 Å), maintaining capsid symmetry and a more compact, less solvent-accessible structure. In this state, the SASA is reduced (∼350,000 Å^2^), suggesting that this empty capsid is less permeable to solvent infiltration compared to № 1. Furthermore, the decomposed r_gyr_ components remain nearly identical. Although a few monomers (e.g., C22 and C31) exhibit higher flexibility, the overall RMSF distribution is relatively narrow, indicating a homogenous dynamic behavior under these conditions. In contrast, for the nucleocapsid under physiological salt conditions (№ 3), the presence of only physiological NaCl led to ongoing structural reorganization and dynamic heterogeneity across subunits. Notably, № 3 exhibited RMSD fluctuations of ∼ 2 Å between consecutive nanosecond intervals, indicating an assembly that remains dynamically unstable. The r_gyr_ further reflected these fluctuations. After a dramatic drop at 40 ns due to capsid elongation, r_gyr_ increased, which is attributed to ongoing capsid reorganization. Moreover, the capsid SASA in № 3 is the highest among all conditions, reflecting the formation of large collective reorganization exposed to the solvent. The broad distribution of monomeric RMSF values further confirms a system-wide dynamic response of the nucleocapsid under physiological conditions (Fig. S6). To ensure the reproducibility of these observations, the conventional MD simulations for assemblies № 1–3 were additionally duplicated. The resulting averaged structural metrics demonstrated good consistency across replicas, confirming that the observed dynamical trends are intrinsic rather than stochastic. The corresponding RMSD and r_gyr_ time profiles for all replicas, along with standard deviation, are provided in the SI (Fig.S1-S3).

The critical role of divalent ions in stabilizing RNA structural integrity has been extensively documented^69–72^. These ions are integral components of certain viral proteins and play a pivotal role in their survival and pathogenicity^73^, though the precise mechanisms remain to be fully elucidated. In our study, we further examined three nucleocapsid conditions (№ 4–№ 6) by varying concentrations of NaCl while including Mg^2+^ to neutralize the RNA, as suggested by previous studies^20^. Interestingly, the nucleocapsid in the neutralized conditions (№ 4) exhibited a relatively slow and uniform increase in RMSD, r_gyr_, and its principal components, as well as SASA, accompanied by a narrow RMSF distribution, where only a few monomers exhibiting higher fluctuations while the majority remain stable and relatively low (of ∼ 3 Å). In contrast, the addition of physiological salt concentration (№ 5), led to larger RMSD oscillations (for up to 60 ns) before gradually stabilizing, suggesting that the virus may require an extended adaptation period under neutral physiological conditions. Under these conditions, the RMSF distribution was skewed toward lower values, and the r_gyr_ principal components remained largely uniform, with only a slight elongation along the *z*-axis. Under more acidic conditions with elevated salt concentration (№ 6), the RMSD and RMSF profiles were comparable to those observed in № 5. However, the global r_gyr_ was reduced over time, resulting in a total capsid compression (at ∼ 70 ns). In this state, the principal axes of r_gyr_ were uniformly distributed and equidistant (Fig. S5), and the capsid SASA decreased, indicating a surface that becomes less accessible to solvent infiltration.

The observed global dynamical signatures between neutral and acidic environments could be linked to changes in inter-monomer electrostatic interactions, particularly salt bridges, which are known to contribute to capsid stability and flexibility. Comparative analysis of salt-bridge networks revealed that assemblies at pH=7 featured a relatively high number of salt-bridge patterns that consistently occur across symmetric monomer pairs. In contrast, under acidic conditions, the salt bridge distribution is markedly reorganized, with only three distinct residue pairs engaging a far broader range of consecutive monomer pairs (see Tables S1-S2 and Fig. S4 for details).

Taken together, the ensemble of MD observations indicates that the STMV capsid exhibits large-scale, concerted “breathing” motions—quasi-elastic expansions and contractions that continuously redistribute structural strain across monomers without loss of overall icosahedral organization. This intrinsic flexibility allows the capsid to dynamically respond to environmental cues, including ionic composition and pH, likely preparing it for functional adaptation while preserving the integrity of the viral shell.

### RNA Dynamics Within Capsid

To further elucidate the interplay between the encapsidated RNA and the capsid dynamics, we investigated RNA behavior along our conventional simulations. Specifically, we analyzed RNA RMSD, r_gyr_, radial distribution function (g(r)), and SASA across nucleocapsid-related STMV conditions (№ 3-№ 6, Table 1) (Fig. 3). The strong similarity in overall curve shape between these RNA profiles and those observed for the capsid (see also Fig. 2A,C,D) indicates a tight dynamic coupling between RNA and capsid motions.

**Figure 3.**
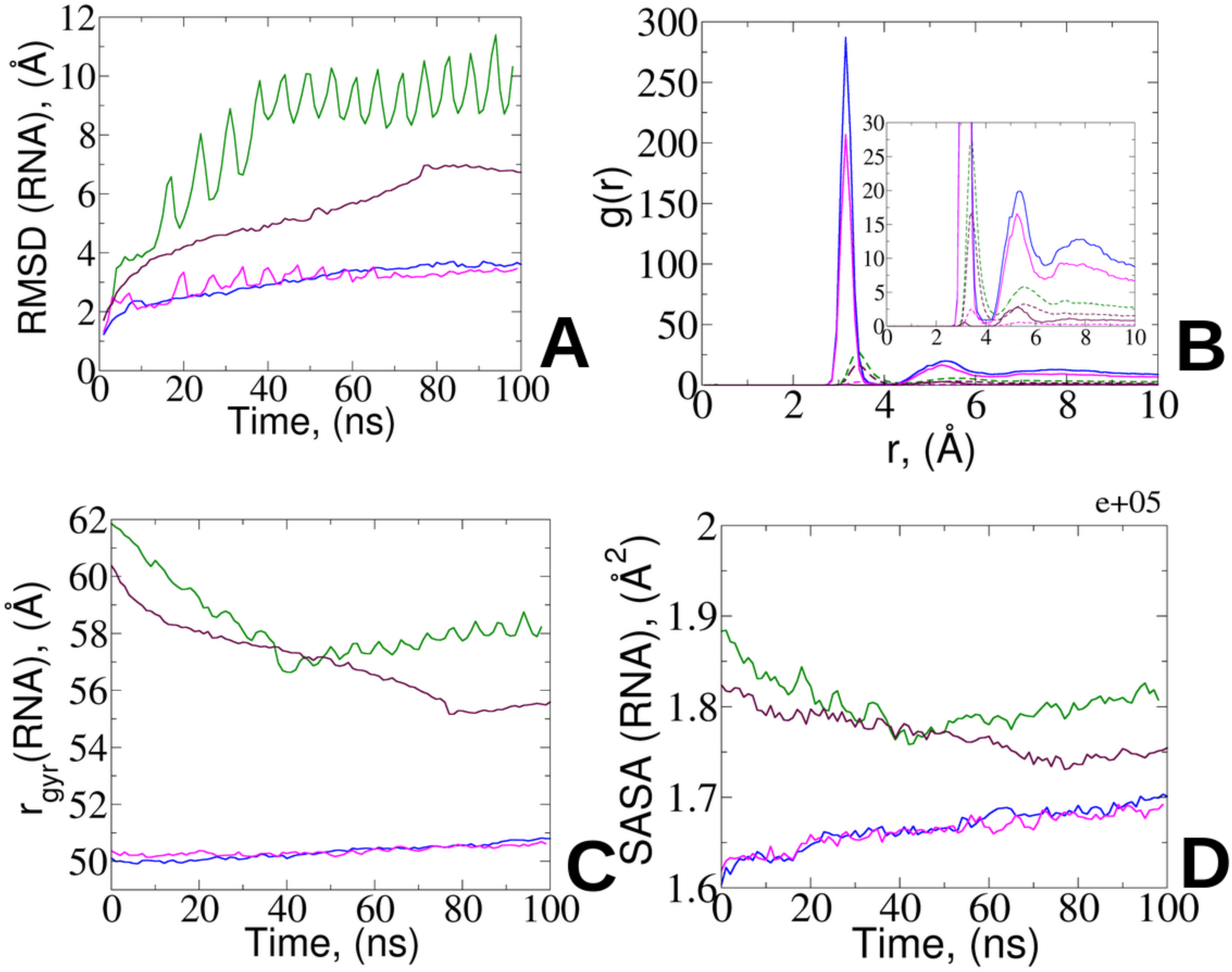
Analysis of the STMV RNA structural dynamics along conventional 100-ns-MD simulations: (A) Root Mean Square Deviation (RMSD) of nucleotide phosphorus atoms, (B) Normalized radial distribution function (RDF) between phosphorous atoms and Mg^2+^ (solid lines) and Na^+^ (dashed lines), (C) RNA radius of gyration (r_gyr_), (D) RNA SASA across the different STMV conditions. The corresponding colors for each studied viral conditions are: № 3 - nucleocapsid in physiological NaCl concentration (green), № 4 - nucleocapsid neutralized with MgCl_2_ (blue), № 5 - nucleocapsid in physiological NaCl concentration with Mg^2+^ present (magenta), and № 6 - nucle- ocapsid in 0.5 M NaCl concentration with Mg^2+^ mimicking pH=5 conditions (brown). Persistent oscillations confirm non-stationary collective rearrangements.

Precisely, the RMSD analysis (Fig. 3A) reveals that, with the exception of condition № 6 (pH=5), the RNA RMSD values closely match those of their corresponding capsids (Fig. 2A), reflecting coordinated structural fluctuations and joint stabilization under neutral pH. Under acidic environment, where protonation predominantly alters the capsid while the RNA retains its native charge state, we observed a significant increase in RNA conformational dynamics. Such enhanced flexibility is likely attributed to the disruption and reorganization of electrostatic interactions as a consequence of capsid protonation. Supporting these findings, the analysis of the radius of gyration (Fig. 3C) further underscores the sensitivity of RNA compaction to both ionic conditions and pH. At pH=7 under physiological concentration with Mg^2+^ present (№ 5), as well as in the presence of MgCl_2_ alone (№ 4), the RNA structure remains comparably compact (∼50 Å). Conversely, under purely physiological conditions (№ 3) and in an environment emulating the acidic milieu (№ 6), the RNA undergoes expansion, adopting a more solvent-accessible conformation (Fig. 3D), highlighting the critical interplay between pH and ionic composition in directing RNA folding dynamics within the capsid.

At this point, we hypothesized that protonation- and ion-induced reorganization of electrostatic networks engages the arginine-rich motifs (ARMs), key mediators of RNA binding in many nucleocapsids, to dynamically adjust RNA–protein interactions under varying environmental conditions^74–76^. Specifically, ARMs are essential in establishing a set of salt bridges with the phosphate of RNA, which can be dynamically rearranged in adaptation response to pH and ion concentration. As discussed previously (see Table S1-S2 and Fig. S4), the reorganization of capsid salt-bridge networks under acidic conditions leads to notable alterations in the symmetry of intermonomer contacts. Such reorganization coincides with an increased propensity for interactions between positively charged capsid residues, particularly the *ε*-amino group of lysine (LYS) and the guanidinium group of arginine (ARG), and the oxygen atoms of the RNA phosphate groups (Fig. S7–S8). Moreover, while hydrogen bonds between capsid amino groups and RNA phosphate moieties account for roughly 40% of total hydrogen-bond contacts (Fig. S7), the salt bridging between the phosphate and the ARM-responsive residues of ARG and LYS might induce the structural reorganization in response to the pH and ionic strength (Fig. S8). In this context, ARMs might function as pivotal molecular switches that finely modulate capsid-RNA interactions, therefore, fine-tune the structural rearrangements necessary for the biological adaptation of the nucleocapsid. Additional insight into the RNA local environment is provided by the radial distribution function g(r) (Fig. 3C). At pH=7, a well-defined peak at 3.2 Å indicates strong coordination of RNA with Mg^2+^, consistent with the established stabilizing role of divalent ions in maintaining RNA tertiary structure^69,70,77–83^. Under acidic conditions, however, the broadening and attenuation of these peaks suggest more transient and weaker Mg^2+^–RNA interactions. This observation points to a redistribution of ion-mediated stabilization, where divalent coordination diminishes and monovalent species, particularly Na^+^, increasingly contribute to RNA charge screening. Capsid protonation within each capsid subunit at pH=5 further modifies the local electrostatic potential, weakening attractive interactions between the RNA and the arginine-rich motifs (ARMs). Detailed ion density analyses supporting this transition are presented in Fig. S14 and discussed further below.

### Investigation By Means Of Enhanced Sampling

The collective radial fluctuations captured within the 100-ns conventional MD trajectories (Fig. 2C) represent the early relaxation dynamics of the STMV capsid, revealing the onset of large-scale conformational rearrangements. While these simulations effectively resolve the initial structural responses, accessing longer-timescale transitions and equilibrium conformational ensembles requires enhanced sampling strategies.

#### Exploratory Gaussian Accelerated Molecular Dynamics Gaussian Accelerated Molecular Dynamics

To extend sampling beyond the initial relaxation regime, we performed exploratory Gaussian accelerated MD (GaMD) simulations on assembly № 2, employing a dual-boost scheme acting on both solute and solvent dihedral and total potential energies^60^. As demonstrated by Miao et al.^57,58^, GaMD typically accelerates conformational sampling by one to two orders of magnitude. In this context, the performed GaMD trajectory corresponds approximately to the conformational sampling expected from about a *µ*s of conventional MD. It should be noted that accessing the millisecond-timescale is currently out of reach for AMOEBA, even for GAMD simulations as it would required to perform around a *µ*s of GAMD simulation which would require several years of computation even when using multiple GPUs. Analysis of the GaMD boost potentials (Fig. S9–S10) confirmed stable, near-Gaussian distributions for both dihedral and total potential boosts, verifying the reliability of the dual-boost protocol. During GaMD, the STMV capsid continued to exhibit slow, collective structural fluctuations similar to those observed in conventional MD (see Fig. S11). The radius of gyration oscillated around 69.5 Å, corresponding to the minimum of the potential-of-mean-force (PMF) profile discussed below (Fig. 4). These fluctuations did not converge to a fixed plateau within the accessible time window, suggesting that full structural relaxation likely proceeds on longer timescales^56^.

**Figure 4.**
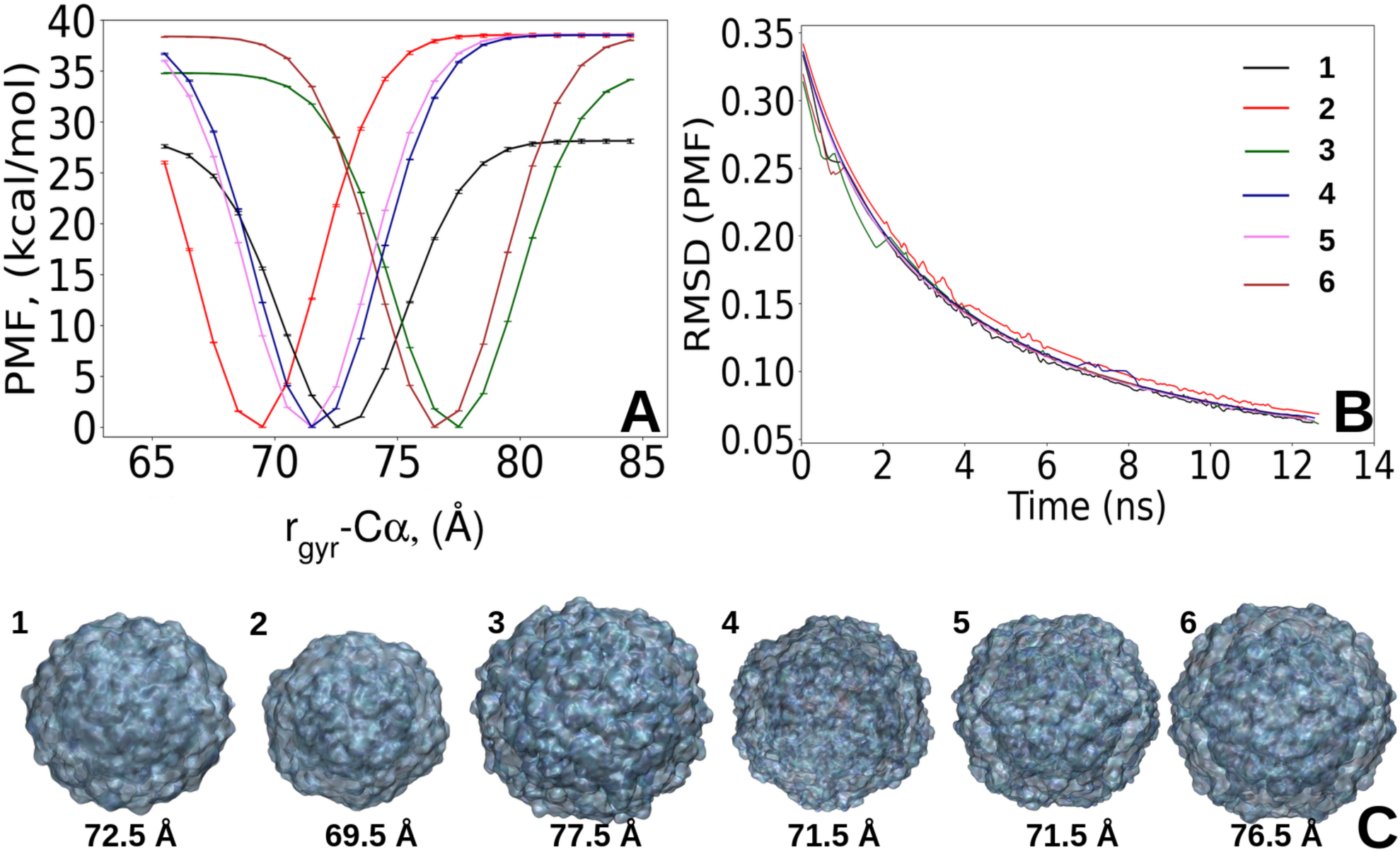
(A) Averaged r_gyr_ PMF with corresponding error bars and (B) averaged convergence plot of r_gyr_ PMF from four independent replicas, with color codes matching the studied conditions (identical to Fig. 2). (C) The capsid structures corresponding to the energy minima of the PMFs.

Although neither conventional MD nor GaMD produced a single, time-invariant structural plateau for the investigated STMV assemblies, these trajectories remain essential to characterizing the assembly dynamical response and the early relaxation pathways sampled on the fully polarizable atomistic potential. WTMetaD was therefore used as a complementary, targeted approach to quantify the free-energy structure along a physically meaningful coordinate (the capsid radius of gyration, r_gyr_). Crucially, the PMFs reported below are converged with respect to this CV and thus reliably identify locally accessible free-energy basins for the capsid radius under the simulated conditions. They should not, however, be interpreted as a map of global equilibrium across the full, high-dimensional conformational space: WTMetaD defines free energies projected onto the chosen CV, while conventional MD and GaMD sample orthogonal slow modes that are not fully collapsed onto r_gyr_.

#### Potential Mean Force Calculations

To clarify the thermodynamic basis of the observed behavior and to identify the dominant conformational basins populated under the examined conditions, WTMetaD^63–65^ simulations were subsequently employed to reconstruct the free-energy landscape along r_gyr_ of the capsid shell defined as the collective variable (CV). Remarkably, the resulting PMF profiles, generated from four independent WTMetaD replicas, exhibited well-defined and highly converged minima (Fig. 4A,B, S12). The small variance among replicas and rapid convergence indicate that each system sampled a consistent, stable free-energy basin, corresponding to the locally accessible thermodynamic states of the capsid.

Specifically, the comparative analysis of PMF profiles revealed distinct energetic signatures depending on RNA encapsidation and environmental conditions, highlighting the intricate coupling between electrostatics and capsid mechanics. Among the RNA-free capsids, № 2 exhibited one of the narrowest minima, centered at 69.5 Å (Fig. 4C), reflecting a compact and energetically stable conformation with limited propensity for expansion. In contrast, № 1 showed a broader PMF minimum and a wider distribution of sampled radii (see Fig. S7), centered at 72.5 Å, indicative of increased structural flexibility and a higher likelihood of empty capsid expansion. The observed difference highlights the critical influence of ionic screening in modulating capsid mechanical properties and underscores their importance in maintaining structural cohesion of empty viral shells in the absence of genome.

Among the RNA-encapsidated assemblies, the PMF landscapes further emphasized the interplay between ion composition and environmental pH. Assembly № 3 exhibited the largest gyration radius, expanding to 77.5 Å, quantitatively confirming that removal of divalent ions under neutral pH promotes capsid expansion through uncompensated electrostatic repulsion between capsid and RNA. Assemblies № 4 and № 5, both containing RNA and Mg^2+^ under neutral pH, converged to nearly identical PMF minima, demonstrating that Mg^2+^ coordination under neutral pH stabilizes a compact conformational basin largely insensitive to bulk salt concentration. Assembly № 6, representing acidic and high-salt conditions, reached a PMF minimum at 76.5 Å, favoring expansion and reflecting protonation-induced weakening of interfacial screening. Collectively, these PMF profiles quantify how subtle shifts in ionic environment translate into global mechanical responses of the viral shell.

### Water and Ion Flux into the Capsid

The examination of conventional MD trajectories additionally revealed measurable solvent and ion permeation through transient inter-monomer gaps (i.e., voids) (Fig. 2), indicating that observed structural oscillations periodically open nanoscopic pores that mediate exchange between the lumen and the bulk solution. To quantitatively assess gap formation and capture structural transitions under varying conditions, we classified interactions between monomers into two distinct categories: stable, tightly packed associations (”occluded pairs”) and transient, more open interfaces (”pore-forming pairs”), as depicted in Fig. S13. Notably, among the empty capsids, № 2 exhibited a more compact inter-monomer arrangement compared to № 1, consistent with our observations on the capsid dynamics (Fig. 2C). For the nucleocapsid-related assemblies, № 3 displayed the fewest number of occluded and pore-forming pairs, yet exhibited the highest solvent accessibility. Although the number of occluded pairs remains largely conserved across the assemblies, the composition and dynamic behavior of the pore-forming interfaces differ, underscoring that ionic strength and pH might influence the kinetics of interface opening. In the nucleocapsid neutralized with MgCl_2_ (№ 4), the pore-forming clusters were less exposed to the solvent compared to № 3, enabling moderate solvent penetration into the capsid interior. Furthermore, № 5 and № 6, both containing Mg^2+^ and Na^+^, demonstrated divergent pore-forming characteristics. In № 5, pore-forming interfaces fluctuated more markedly in solvent accessibility, whereas in № 6, this parameter was considerably more stable, suggesting that elevated ionic strength and acidic pH conditions in № 6 may enhance inter-monomer stability while still permitting controlled pore formation.

To further elucidate the spatial characteristics of solvent permeability, we quantified the concentration ratio of water molecules inside the capsid relative to the bulk environment (Fig. 5A). In addition, radial distribution analyses were performed enabling a consistent comparative assessment of solvent localization across all assemblies (Fig. 5B). The most prominent water accumulation, along with a progressive water influx, was observed for the empty capsid within physiological concentration (№ 2), indicating that confined solvent might participate in electrostatic relaxation and support capsid contraction/expansion dynamics without compromising the shell geometry. While nucleocapsid № 3 demonstrates the most striking water influx, indicative of its heightened dynamic instability (Fig. 2A), its internal water concentration ratio is notably lower than that observed for № 2. Among the rest, assemblies № 5 and № 6 exhibit similar solvent permeability profiles, whereas № 4 shows the minimal degree of water influx.

**Figure 5.**
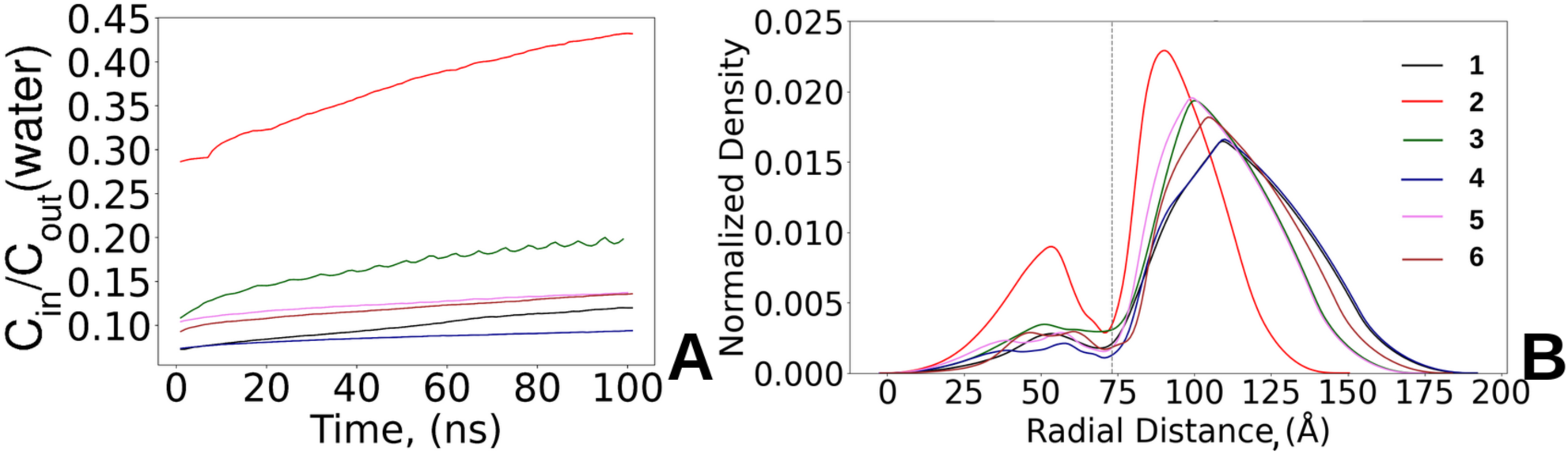
Water permeability along different conditions. (A) Concentration ratio of water molecules inside the capsid to the water molecules in the bulk. (B) Normalized radial distribution of water molecules. The dashed line corresponds to the averaged capsid gyration radius across all assemblies. From the left side from the dashed line are the water radial distribution within capsid boundaries, from the right – water radial distribution within the bulk.

The decoupling between pronounced inter-monomer pore dynamics and relatively low net water permeability indicates that these openings might primarily facilitate ion exchange rather than bulk solvent flow. To probe this, we quantified net ion fluxes into both the capsid (Fig. 6A–F) and RNA-containing interiors (Fig. S15), and analyzed radial ion density distributions (Fig. S14) to evaluate preferential ion accumulation.

**Figure 6.**
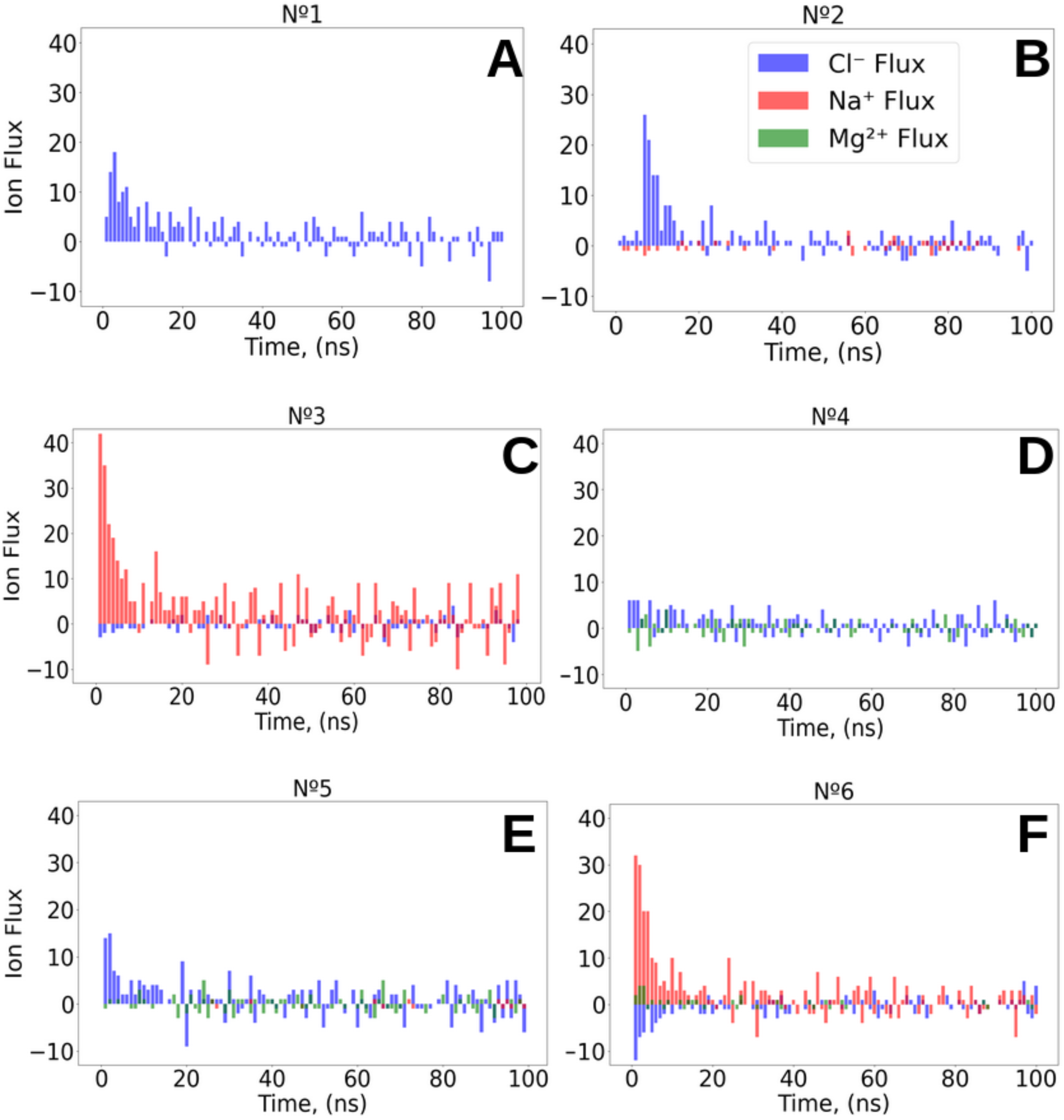
Net Ion flux of Cl^−^ , Na^+^, and Mg^2+^ displayed in blue, red, and green, respectively for each of the studied STMV assemblies of № 1–6 represented accordingly in A–F plots. The negative values of flux correspond to the ionic efflux from the capsid.

In empty capsids, Cl^−^ ions are preferentially accumulated within the interior cavity, exhibiting a broader distribution in № 1 than in № 2, consistent with weaker electrostatic confinement at lower salt concentration. Thus, Na^+^ in № 2 tend to remain external to the capsid, effectively counterbalancing the excess Cl^−^ . Within the nucleocapsids, RNA introduces strong local electrostatic gradients that attract counterions toward phosphate moieties. In № 3, monovalent Na^+^ ions penetrate into the RNA-rich core, neutralizing backbone charge and replacing Cl^−^ as primary internal species.

Assemblies № 4 and № 5 exhibit similar ionic profiles, marked by preferential accumulation of Mg^2+^ near RNA and Cl^−^ at capsid periphery. In № 6, although the water permeability profile resembles that of № 5, the ionic influx is dominated by Na^+^ rather than Mg^2+^. At pH=5, protonation of acidic residues electrostatically exclude Cl^−^ ions and promote Na^+^ influx. This behavior is reminiscent of a Donnan effect, where fixed charges in a confined volume drive ion redistribution and modulate the electrostatic environment. Together, these observations indicate that RNA influences capsid permeability not only through structural confinement but also via electrostatic interactions, likely tuning RNA flexibility and overall capsid dynamics.

### Residence Time

To elucidate the ionic influence on capsid-RNA interactions, we analyzed the preferential ion binding sites and quantified the residence times of interactions between ions and charged residues within the capsid and RNA. At physiological pH, neither Na^+^ nor Mg^2+^ exhibited interactions with the negatively charged side chains of capsid residues (i.e., ASP and GLU), suggesting that for assemblies of № 1–5, these cations do not directly mediate inter-monomer interactions. Instead, their interactions are predominantly confined to the phosphate backbone of RNA, consistent with dynamic, transient ion-phosphate coordination. It was already mentioned above that a distinct behavior was observed in № 6 at acidic pH. Herein, Na^+^ ions localized to the negatively charged carboxylate oxygens of GLU residues (see Fig. S14–S15), corresponding to an initial influx of Na^+^ ions at the beginning of the simulation (Fig. S17), where a small subpopulation (∼24 ions, ∼ 1% of the total Na^+^) formed long-residence interactions with GLU residues. These interactions participated in dynamic ion exchange within the capsid environment, facilitating cationic entry into the cavity and their subsequent engagement with RNA phosphate groups (Fig. 7D). Although less abundant, Mg^2+^ (∼4 ions) displayed pronounced long-residence interactions with the RNA phosphate groups, underscoring the genome structural adaptability under acidic conditions. Across other nucleocapsid assemblies at pH=7, both cations participated in dynamic RNA interactions, Na^+^ favoring short-lived contacts, and Mg^2+^ exhibiting stronger, longer-lasting associations.

**Figure 7.**
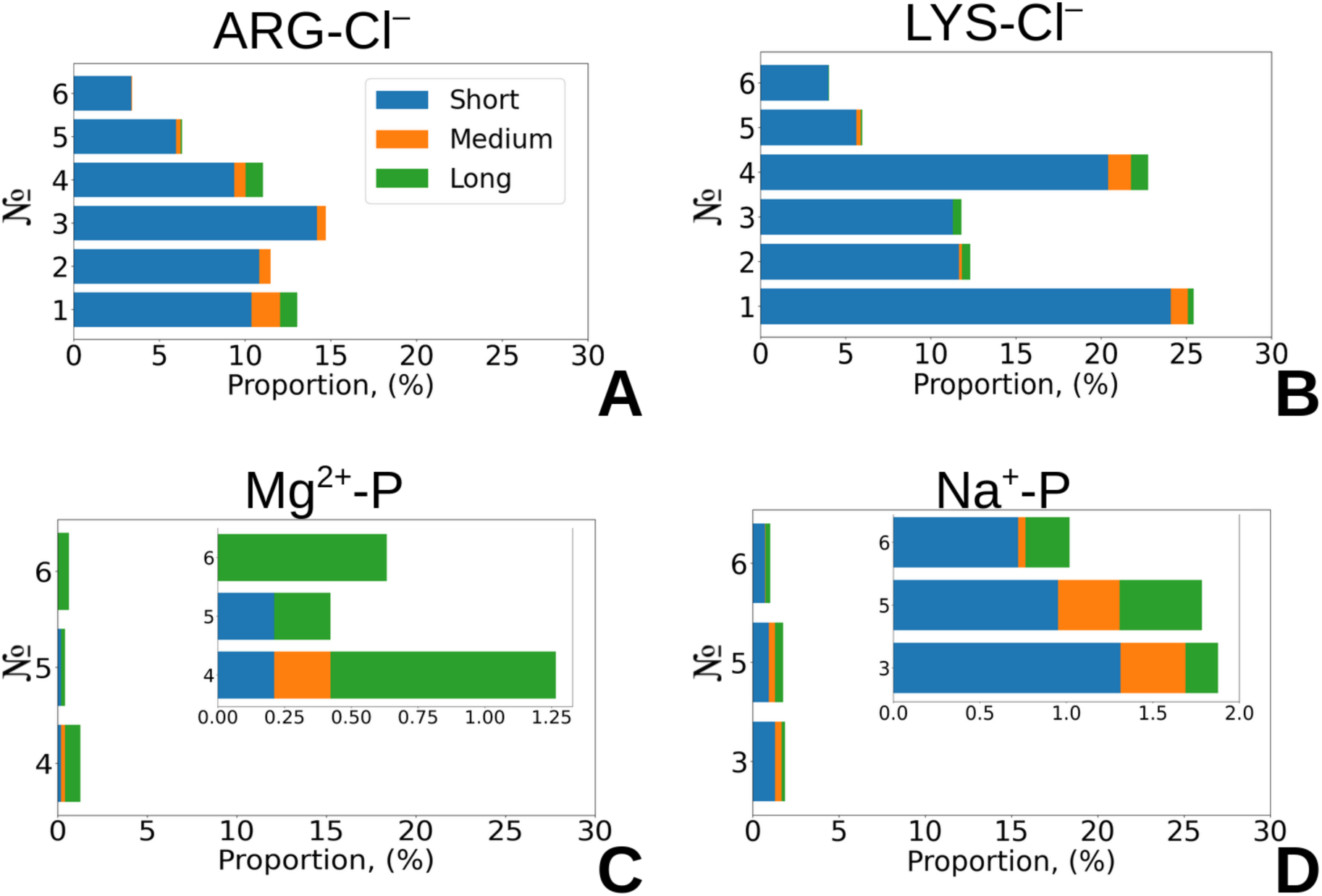
Residence time stacked bar charts for Cl^−^ ions interacting with positively charged residues in six STMV assemblies. (A) Side-chain ARG interactions with Cl^−^ . (B) Side-chain LYS interactions with Cl^−^ . Phosphorus atom of phosphate group of RNA interactions with (C) Mg^2+^ and (D) with Na^+^. Bars represent the proportions of ions that observed within the residence times categorized as Short (blue, 1–3 ns), Medium (orange, 4–7 ns), and Long (green, *>*7 ns). *X*-axis (Proportion,(%) indicates the fraction of ions in each residence-time category. STMV assembly labels (№ 1–6) correspond to different simulation conditions, as described in Table 1.

Besides, Cl^−^ exhibits predominantly short-residence interactions with positively charged ARM residues (i.e., ARG, LYS), transiently neutralizing electrostatic repulsion and actively destabilizing existing inter-monomer salt bridges. This behavior effectively promotes the formation of transient pores (Fig. 7A-D), highlighting Cl^−^ as a key modulator of capsid structural plasticity.

### The Role of Ions in Inter-Monomer Salt-Bridge Disruption

The pronounced influence of Cl^−^ on ARM-mediated contacts motivated a systematic investigation into how specific ions modulate inter-monomer associations and regulate capsid dynamics. To this end, we performed a comprehensive analysis of salt-bridge disruption and evaluated the role of polarizability in this process, with № 1 serving as the system for displaying our observations. Qualitatively identical behavior was observed in all other STMV conditions, underscoring the generality of the mechanism. As a representative example, the C4-C5 dimer initially hosts two closely apposed salt-bridge triplets at pH = 7 (Fig. S4): namely, 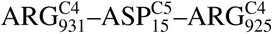 and 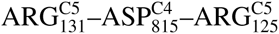 (Fig. 8A,B).

**Figure 8.**
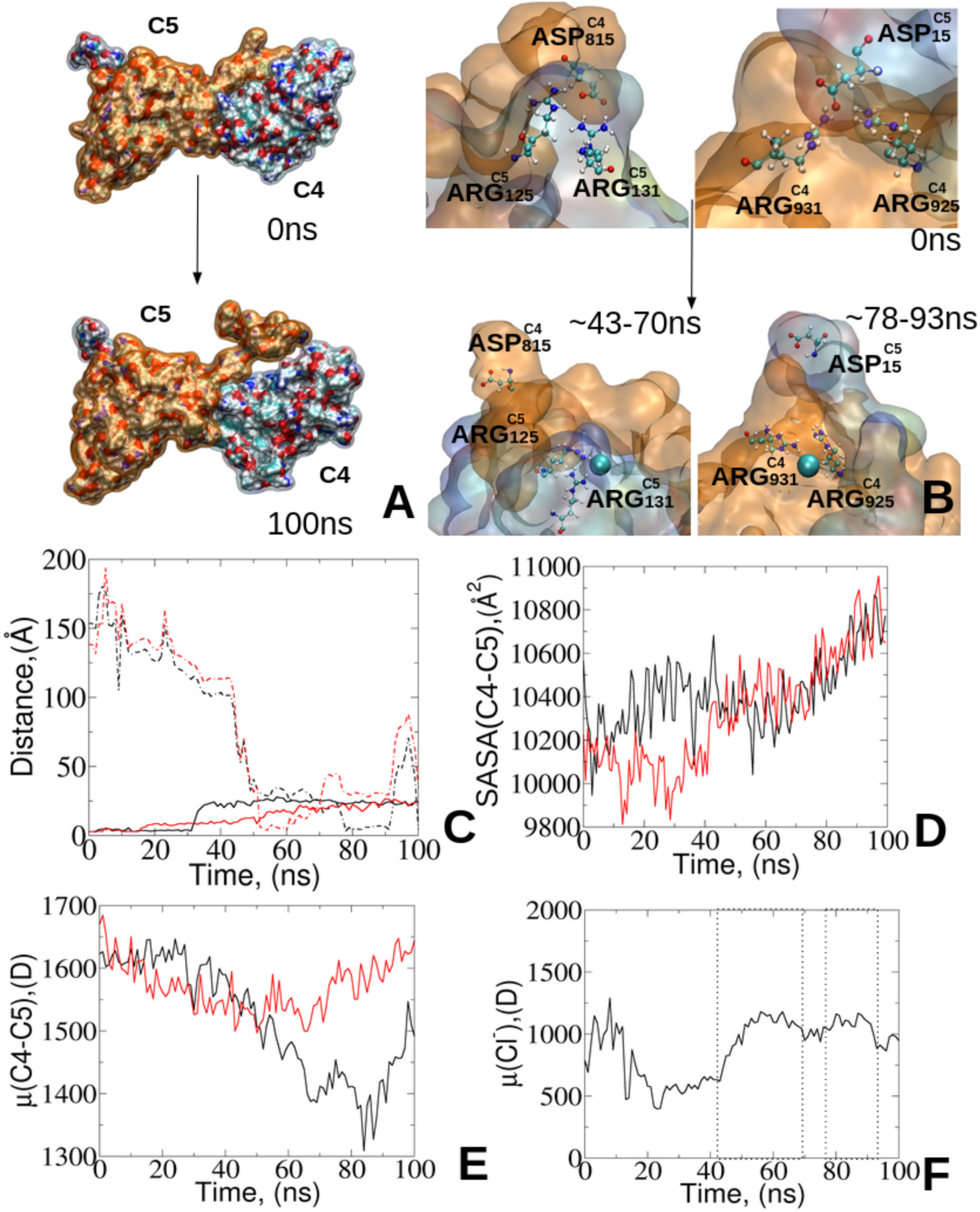
(A) Representation of the disruption of salt bridges between the C4 (orange) and C5 (blue) monomers, showing the initial capsid configuration (top) and the final 100-ns simulation frame (bottom). (B) Zoom-in view of the salt-bridge triplets of 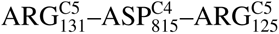 and 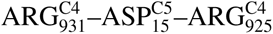 (top) and their disruption along the trajectory by the same Cl^−^ ion shown in cyan van der Waals representation. (C-F) Time-dependent analysis of the salt-bridge disruption between C4 and C5 monomers in the neutralized capsid (№ 1) by Cl^−^ . (C) Distance between the charged residues: 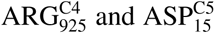 (solid black), the Cl^−^ ion approaching 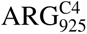 (dashed black), and 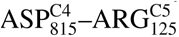 (solid red) and Cl^−^ ion approaching 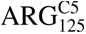 (dashed red). (D) Solvent Accessible Surface Area (SASA) of the C4 (black) and C5 (red) monomers. (E) Induced dipole moment evolution of the C4 (black) and C5 (red) monomers, and (F) of the Cl^−^ ion with vertical lines corresponding to the periods of time, where Cl^−^ disrupts 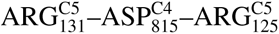 and 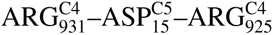.

During the 100-ns simulation, a transient pore opening event was observed at the C4-C5 interface (Fig. S13), coincided with solvent exposure of both monomers (Fig. 8D). This event is initiated by the binding of Cl^−^ to the interfacial side chains of 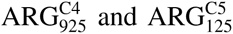 (Fig. 8B), triggering both local rearrangement and global monomer separation.

Between ∼ 43 and 70 ns, a single Cl^−^ ion approaches and binds 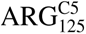, reducing their interatomic distance (Fig. 8C) and effectively saturating the positive site on C5 monomer that precludes the reformation of the original salt bridge (Fig. 8D). Concurrently, the ion induced dipole moment increases sharply, indicating its transition from bulk solvation to direct protein contact (Fig. 8F). Ion binding drives local rearrangements of 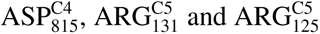, reorienting them toward solvent, as confirmed by their elevated dipole moments (Fig. S18).

Over the next ∼8 ns, the same Cl^−^ remains close to the C4-C5 interface, and begins to engage 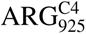, further weakening the 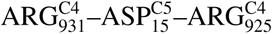. In response, 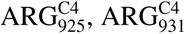 and 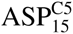 ASP^C5^ rotate inward toward the protein core, each exhibiting a decrease in induced dipole. At the monomer level, these changes manifest as a net dipole-moment decrease in C4 and an increase in C5 (Fig. 8E), representing a decisive step in destabilizing the C4–C5 contact.

Together, these observations demonstrate that transient Cl^−^ binding actively destabilizes persistent salt bridges and induces local dipole realignment, triggering reversible pore openings, thus, providing a molecular mechanism for capsid plasticity and controlled permeability.

### Conclusion and Perspectives

Performing a cumulated total of 1.5-*µ*s of simulation using a polarizable force field, including 1.2-*mu*s of conventional MD combined with additional dual-boost GaMD WTMetaD simulations, we revealed that the STMV capsid behaves as an electrostatically responsive and dynamically adaptive assembly. Its conformational behavior emerges from a fine balance among protein–protein salt bridges, RNA-protein charge compensation, and ion-mediated electrostatic screening.

At the heart of this adaptability lies a dual electrostatic mechanism. Under neutral, physiological conditions, divalent cations (Mg^2+^) bridge the RNA phosphate backbone and the arginine-rich motifs (ARMs) within the capsid, acting as structural crosslinkers that compact the genome and rigidify the protein lattice. This coupling minimizes internal charge repulsion, stabilizing the virus in a compact, low-energy configuration. Conversely, transient binding of monovalent ions, particularly Cl^−^ , locally destabilizes persistent salt bridges between subunits. These short-lived events realign local dipole moments and generate nanoscopic pores that mediate selective water and ion exchange. The interplay between these two regimes, crosslinking and disruption, produces the reversible quasi-elastic dynamics of the protein shell that define STMV structural plasticity. Besides, our atomistic trajectories also demonstrate that even in the absence of RNA, the capsid can persist as a near-native, icosahedral structure while allowing substantial conformational flexibility especially when environmental screening is sufficient (for example, under physiological NaCl).

Under acidic and high-ionic-strength environments, protonation of acidic residues reorganizes the interfacial charge network and partially redistributes cations within the capsid interior. This adjustment weakens divalent coordination (i.e., Mg^2+^) while enhancing monovalent mobility (i.e., Na^+^), leading to a controlled expansion of the viral shell. The ability to reversibly switch between compact and expanded conformations through subtle changes in protonation and ionic composition underscores STMV environmental sensing capability, a hallmark of its biophysical adaptability.

While our investigation target preassembled capsids rather than full assembly or disassembly, they offer atomic-level insight into the electrostatic and mechanical couplings that define STMV behavior at its early pathways, including RNA encapsidation, ion homeostasis, and controlled permeability during intracellular trafficking. From a broader mechanistic standpoint, STMV exemplifies how fine-tuned electrostatic cooperativity may govern a mechanical resilience in related ssRNA viruses. Experimental validation will be further essential to refine this mechanistic framework and to elucidate how ion-regulated dynamics contribute to viral function and infectivity.

## Methods

### Molecular Dynamics Protocol

Details regarding the composition of the studied STMV conditions are provided in Table 1. The configuration files, based on PDB ID of 1A34^5^, corresponding to the studied systems № 1 and № 4 were obtained from the Urbana-Champaign Theoretical and Computational Biophysics Group website^84^. For all other systems studied at pH=7, the capsid (total of 135,960 atoms) and nucleic acid (total of 30,330 atoms) structures were obtained from these files. The net charge of the protein at pH=7 was +300 and the RNA carried a charge of −949 raising from the negatively charged phosphate backbone. The total net charge of all the systems, including ions and water, was set to neutral within VMD psf/pdb generation workflow together with its built-in solvation and ionization plugins^67^. This procedure placed explicit water molecules and counterions around the viral shell to achieve the target ionic strengths (0.15 M or 0.5 M NaCl, depending on the system) while maintaining overall charge neutrality. For № 6, the capsid protonation at pH=5 was performed using the APBS-PDB2PQR software suite^85^, where the PDB-structure was loaded, and the appropriate protonation states were assigned. Under these conditions, among all charged residues, only GLU_108_ was protonated, making the net charge of the protein at pH=5 of +360, accordingly. The RNA net charge remains the same at pH=7 and pH=5, as it contains only adenine (ADE) and uracil (URA), whose pKa values fall outside this range^86^, requiring no protonation adjustments at pH=5. The input files were then prepared for further manipulations using the CHARMM-GUI open-source platform^87–89^ and VMD^67^ for solvation and ionization. The output PDB-formatted files were then subsequently parameterized with the AMOEBA polarizable force field using the **PDBXYZ** utility available in the TINKER 8 software package^90^. The application of a fully polarizable force field such AMOEBA^47–49^ allows the simulation to reflect the dynamic redistribution of electronic charge in response to the local environment, providing a more accurate and detailed description of intermolecular interactions. Our MD simulations were performed using the Tinker-HP v1.2. Comprehensive documentation of the TINKER source code can be found in references 91,92. Each system was subjected to an initial energy minimization using the limited-memory Broyden–Fletcher–Goldfarb–Shanno (L-BFGS) algorithm, with the minimization process terminating upon achieving a root mean square (RMS) gradient of 1 kcal/mol. Following satisfactory convergence, the systems were prepared for conventional molecular dynamics (MD) simulations, preliminary pre-equilibrated in NVT ensemble. MD production simulations in the NPT ensemble were then conducted using the BAOAB-RESPA1 integrator^93^, implemented within the GPU-accelerated Tinker-HP software package^94,95^, which is publicly available through the GitHub repository^92^ and the Tinker-HP website^96^. The integration scheme utilized three distinct time steps: a short time step of 0.001 ps for accurately resolving the fastest motions within the system, an intermediate time step of 0.333 ps to capture medium-frequency motions and interactions efficiently and a 10-fs time step for slowly varying long-range interactions. Given the large size of our systems, exceeding 1 million atoms, each simulation was performed utilizing 8 H100 GPUs, highlighting the exceptional scalability of the Tinker-HP code.

### Gaussian Accelerated Molecular Dynamics Simulations

To enhance conformational sampling, we employed Gaussian accelerated molecular dynamics (GaMD)^57,58,60^ as implemented in the Tinker-HP package^94,95^. GaMD accelerates barrier-crossing events by adding a smoothly varying boost potential to the system energy surfaces, thereby reducing energy barriers while maintaining accurate Boltzmann-weighted statistics.

The GaMD protocol consisted of several stages of preparatory and equilibration steps designed to ensure accurate determination of the boost potential parameters. An initial conventional MD equilibration of 10,000 steps was followed by 200,000 conventional MD sampling steps to collect potential energy statistics. These data were used to parameterize the GaMD boost potential. Subsequent equilibration under GaMD biasing included 100,000 preparatory steps and 100,000 production equilibration steps. Both the total potential energy and dihedral energy terms were boosted, enabling enhanced exploration of both global and local conformational degrees of freedom. The standard deviation upper limit for the boost potential was set to 3 kcal/mol, following previously established protocols^58^. Energy and boost statistics were recorded every 1 ps.

### Potential of Mean Force Calculation

To efficiently explore the free-energy landscape of the STMV capsids, we employed well-tempered meta-dynamics (WTMetaD)^63–65^, an enhanced sampling approach designed to overcome the limitations of conventional MD simulations by dynamically modulating the growth of a bias potential applied to a selected collective variable (CV), thus, preventing overestimation of free-energy barriers and ensuring smooth convergence of the free-energy surface. WTMetaD scales the bias height based on the accumulated bias energy, ensuring that the system remains close to thermodynamic equilibrium, preserving realistic pathways during sampling. For the potential of mean force (PMF) calculation, given the large-scale nature of capsid dynamics, we selected the radius of gyration (r_gyr_) as the CV. The choice of r_gyr_ was informed by an initial analysis of conventional MD simulations, where we observed its gradual increase over time without reaching a plateau, making it an effective descriptor of the capsid global structural changes. For the WTMetaD calculations along r_gyr_ we used the C*_α_* atoms of the capsid, as implemented in the Colvars Module^97–99^ in combination with the Tinker-HP software. To ensure accurate and reproducible results, all PMF calculations were performed in four replicas for each STMV assembly, providing robust statistical averaging and minimizing the risk of biases or inconsistencies in the free-energy estimates. In total, 300 ns of accumulated time (12.5 ns per replica) was simulated.

### Root Mean Square Deviation

The Root Mean Square Deviation (RMSD) analysis was conducted using the VMD software^67^. To accurately quantify structural deviations over time, the capsid C*_α_* atoms were aligned to a reference structure extracted from the initial time frame. This alignment step removes any translational and rotational movements that do not reflect genuine conformational changes, allowing for a more precise comparison of the structural evolution. Following alignment, the RMSD was calculated across all simulation frames. This provided insights into how much the structure deviated from the reference over time, indicating the overall stability of the capsid in the various environmental conditions. The results were then plotted to visualize the trends in structural stability. The same procedure was applied to evaluate the RNA dynamics using the phosphorus atoms as a benchmark.

### Root Mean Square Fluctuation

The Root Mean Square Fluctuation (RMSF) of all capsid C*_α_* atoms was performed using the **measure rmsf** utility available in the Visual Molecular Dynamics (VMD) software package^67^. Additionally, we split the calculation of RMSF by monomeric segments to leverage more details of C*_α_* fluctuations. The post-processing of the RMSF data was performed using Python, with the aid of the `NumPy`^100^, `pandas`^101^, and `matplotlib`^102^ libraries. The RMSF dataset was grouped by segment identifiers to compute mean RMSF and standard deviation values for each monomer. These statistical summaries were visualized using bar plots, where mean RMSF values were shown with corresponding error bars representing the standard deviation. Additionally, RMSF value distributions for individual segments were illustrated using histograms, each rendered in a distinct color derived from a colormap.

### Salt-Bridge Analysis

#### Inter-monomer Salt-Bridge Analysis

To investigate the salt-bridge interactions between monomers in the STMV capsid, we identified the initial salt-bridge patterns from the initial frame of MD simulations using VMD. This initial dataset was used as a reference for subsequent analysis. The extracted patterns were systematically processed to identify residue pairs forming inter-monomer salt bridges, focusing on interactions involving segments labeled from C0 to C59 for assemblies under pH=7 and pH=5.

The data were parsed and grouped by amino acid pairs and corresponding segment interactions. Interaction counts were calculated, excluding self-interactions and ensuring symmetry in the segment pair ordering. These counts were then organized into a matrix representing the frequency of salt-bridge formation between segment pairs.

To visualize the inter-monomer salt-bridge network, a heatmap was constructed, illustrating the interaction density between segments. Thus, the heatmap highlighted regions of high interaction frequency, providing insights into the structural organization and stability of the capsid. Variations in the interaction patterns under different conditions were further analyzed to assess the role of ionic interactions in capsid structural integrity.

#### Capsid-RNA Salt-Bridge Analysis

To quantify electrostatic interactions between the positively charged residues of the STMV capsid and the negatively charged phosphate groups of the RNA, a frame-by-frame salt-bridge analysis was conducted over the entire trajectory also in VMD. For each frame in the trajectory, coordinates of the nitrogen atoms of the charged side chains of arginine (ARG), lysine (LYS) and oxygens of the phosphate groups were extracted and pairwise distances were calculated. Salt bridges were defined as interactions occurring within a 3.2 Å cutoff distance. If such a condition was met, the interaction was recorded. Subsequent statistical analysis and visualization were performed in Python. For each salt bridge, distances were aggregated over time, and average interaction distances along with standard deviations were calculated.

### Capsid-RNA Hydrogen Bonding Analysis

To examine the hydrogen bond interactions within nucleotide-present STMV assemblies under varying environmental conditions, we analyzed hydrogen-bond data derived from the trajectory in VMD^67^. Using a distance criterion of 9 Å and an angle threshold of 90*^◦^* for evaluating protein-RNA hydrogen bonding^103–106^, we assessed RNA-capsid interactions across nucleocapsid-related STMV conditions (№ 3-№ 6, Table 1). The total number of hydrogen bonds for each interaction type was extracted for each nanosecond of the trajectory.

To compare the hydrogen bond distributions, we calculated the percentage contribution of each interaction type to the total hydrogen bond count for each system. This approach allowed us to assess the relative abundance of different hydrogen-bond types, independent of system size. The interaction types were visualized in scatter plots, with each system represented on the x-axis and the hydrogen bond percentage on the y-axis.

### Radius of Gyration

To estimate the radius of gyration (r_gyr_) for the capsid and RNA, VMD^67^ was first used to extract the C*α* and phosphorus atom coordinates from all frames of the conventional MD simulations. These coordinates were then processed in Colvars Dashboard plugin^107^ available in VMD.

To assess the shape of the capsid and determine whether it remains spherical or becomes elongated along its principal axes (i.e., x, y, and z), the moments of inertia tensor for each frame of the simulation were calculated. These moments of inertia describe how the mass of the capsid is distributed relative to the center of mass along each of the principal axes. The principal moments of inertia, *I_xx_*, *I_yy_*, and *I_zz_*, were extracted for each frame. To derive the radii of gyration along these axes, the square roots of the eigenvalues were computed. This calculation yields the principal radii of gyration r_gyr_*_,x_*, r_gyr_*_,y_*, and r_gyr_*_,z_*, which characterize the size of the structure along each axis. If all three principals of radii are roughly the same, the capsid is considered to be spherical, whereas if one or two radii are much larger than the others, the capsid is elongated along those axes.

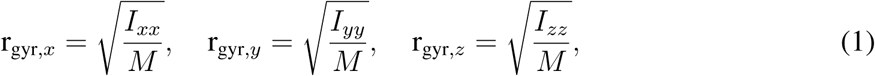

where *M* is the total mass of capsid calculated as a sum of all atomic masses of each atom. The total radius of gyration, *R_gyr_*, is then obtained by combining these components in quadrature:

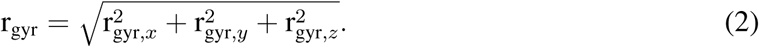

### Water and Ionic Flux And Concentration Ratio Determination

To investigate the permeation of water and ions into the STMV capsid/nucleocapsid and to assess the corresponding water concentration ratio, we used the capsid and RNA radii by determining the simple average distance of all capsid/RNA atoms from their center of mass along the simulation frames using VMD. Under the assumption of spherical geometry, the capsid volume was calculated as:

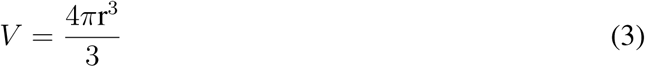

where r represents the computed radius that defined the “inside” region, while the simulation box volume set the framework for the “outside” environment.

Ions and water molecules were then selected, and their radial distances from the capsid center of mass were computed. Species with a radial distance less than the calculated radius were classified as residing “inside” the capsid/RNA, while those with larger radial distances were grouped as “outside.” Complementary histograms of the ion radial distances were generated using the `seaborn` library^108^, where the density profiles were overlaid with a vertical line representing the mean capsid/RNA radius. Normalized density plots were used in the radial density analysis to allow for standardized comparisons of ion distributions independent of the absolute number of each ion type.

Besides, the net flux for each species was then calculated as the difference in the number of particles inside the capsid/RNA between consecutive frames. The flux values for each ion were visualized as separate bar plots over time to illustrate the transient changes in particle counts. Additionally, we performed a computation of water concentration ratio as:

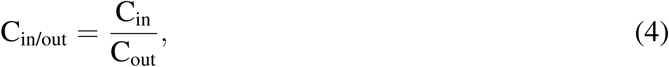

where C_in_ and C_out_ represent the number of water molecules inside and outside the cavity, respectively.

### K-means Clustering

To investigate the capsid porosity, we analyzed the relationship between the center-of-mass (COM) distance and the solvent-accessible surface area (SASA) for monomeric pairs within the capsid. The COM distances were computed using a TCL script in VMD, which calculated the Euclidean distance (via the **vecdist** function) between the COM of each consecutive segment. Similarly, SASA values were determined using a TCL script in VMD, employing **measure sasa** with a probe radius of 1.4 Å to calculate the SASA for each pair. Both COM distance and SASA data were accumulated across all frames of the trajectory. The average and standard deviation for both variables were then computed for each monomeric pair.

To identify meaningful groupings of molecular pairs, K-means clustering was applied to the processed data, with the number of clusters set to four (*k* = 4). The clustering algorithm used both SASA and distance as features, with the aim of partitioning the data into distinct groups based on these properties. Following the clustering, the cluster with the smallest average SASA was identified as the cluster of “occluded monomer pairs,” while the cluster with the largest average SASA was considered the “pore-forming monomer pairs”. These clusters were extracted for detailed analysis. To illustrate the clustering results, a combined scatter plot by means of the `matplotlib` python library^102^ was generated, displaying SASA against COM distance for molecular pairs in both the smallest and largest clusters. Error bars were included to represent the standard deviations of SASA and COM distance, providing insights into the variability of the data. The plot was color-coded, with the occluded pairs cluster represented in blue and the pore-forming cluster in green.

### Ion-Residue Residence Time Analysis

To investigate the dynamics of ion interactions with specific molecular groups, we quantified the residence times of ions near target residues using a TCL script in VMD, employing **measure contacts** within a defined cutoff distance along the trajectory. The cutoff distances used for detecting interactions were derived from the first peak of the radial distribution function (RDF) for each pair of ion and a charged group of a residue side-chain pair, such as LYS (3.2 Å) and ARG (3.4 Å) interacting with Cl^−^ and phosphorus atom of phosphodiester of RNA interacting with Mg^2+^ (3.2 Å) and Na^+^ (3.5 Å) (see Fig. S16 in SI). The residence time for each ion was calculated as the cumulative number of nanoseconds in which the ion remained within the interaction cutoff distance. This allowed for a detailed analysis of ion dynamics, focusing on their short-range interactions.

Building on this analysis, we further grouped the residence time data into three categories—short (1–3 ns), medium (4–7 ns), and long (>7 ns)—to facilitate the interpretation of interaction durations. The total number of ions of each system was used to normalize the residence time counts, allowing meaningful comparisons across systems with varying ion concentrations. The results were visualized using horizontal stacked bar chart plots, where each bar represented the distribution of residence times for a given STMV assembly, residue, and ion type, therefore, highlighting the relative prevalence of short-, medium-, and long-term interactions, providing a clear and comparative view of interaction lifetimes.

## Acknowledgement

This work has received funding from the European Research Council (ERC) under the European Union’s Horizon 2020 research and innovation program (grant agreement No 810367), project EMC2 (JPP). ZV thanks the National Science Foundation (NSF) for funding her research stay (grant CHE-2244028). Computations have been performed at IDRIS (Jean Zay) on GENCI Grants: no A0150712052 (J.-P. P.) and grant GC010815453 (Grand Challenge H100 Jean Zay, J.-P. P.)

## Supporting Information Available

Additional analyses of the trajectories supporting the results of this study.

## Author contribution

**Designed research:** M. B., L. L., J.-P. P. ; **Performed research:** M. B. , Z. V.; **Contributed analytic tools:** O.A ., L. L., J.-P. P.; **Analysis:** M. B., L. L., J.-P. P. ; **Wrote the paper:** M. B., J.-P. P. with the inputs of all authors.

