## Supplementary Information for "Satellite Tobacco Mosaic Virus: Revealing Environmental Drivers of Capsid and Nucleocapsid Plasticity using High-Resolution Simulations"

### Neutralized Capsid (№1) Replicated Simulations

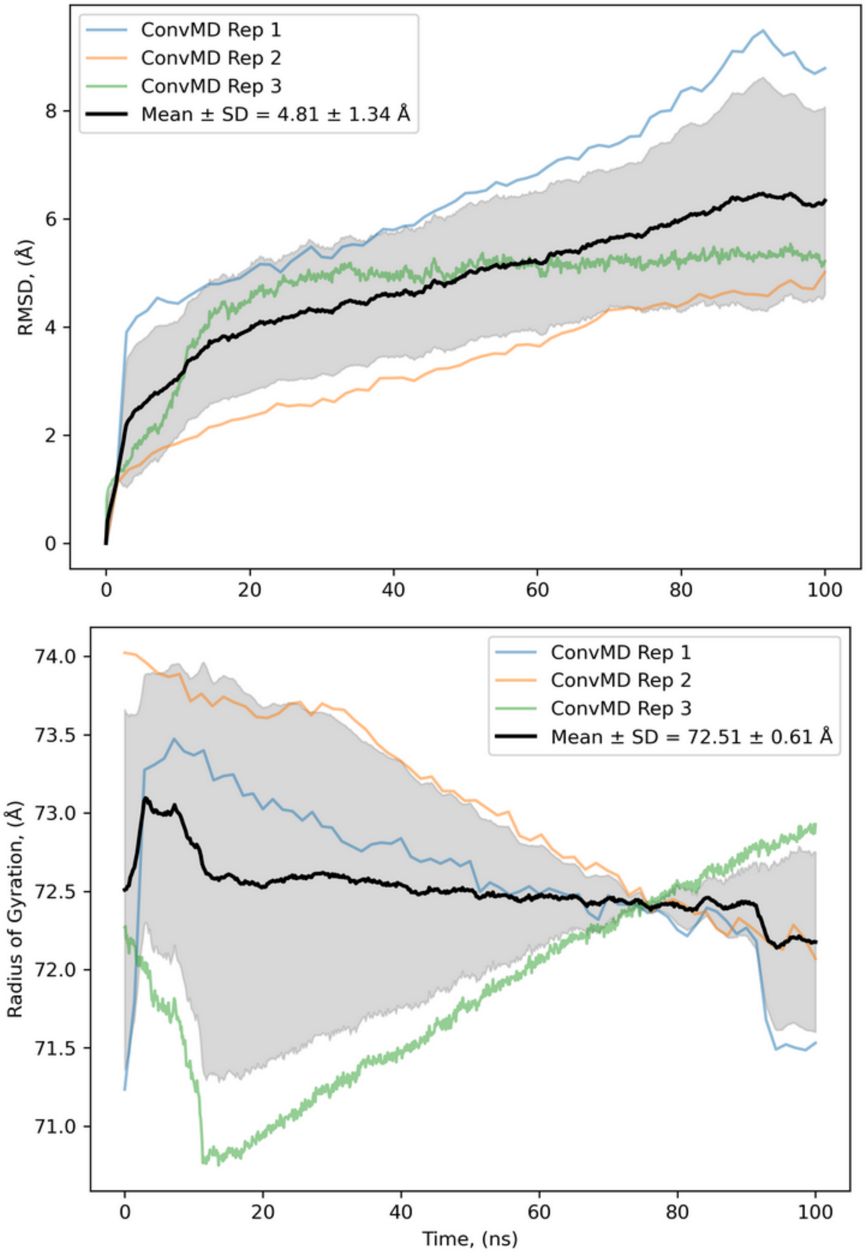


Figure S1. (Top) Root Mean Square Deviation (RMSD) and (bottom) gyration radius of the C$\alpha$ atoms along 100-ns triplicated conventional MD simulation. ConvMD Rep 1 corresponds to the conventional MD trajectory used for analysis in the main text.

### Capsid at Physiological Concentration(№2) Replicated Conventional Simulations

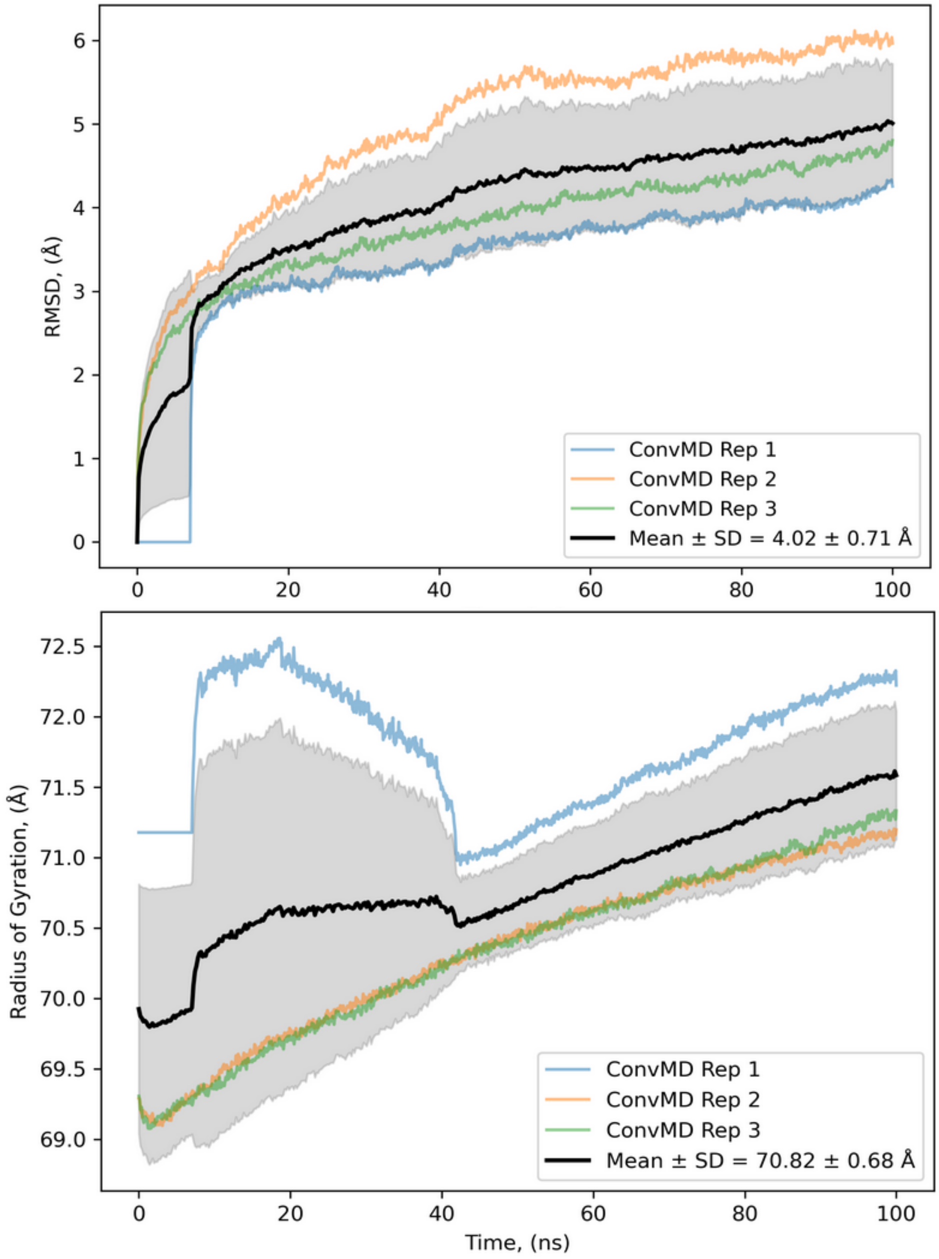


Figure S2. (Top) Root Mean Square Deviation (RMSD) and (bottom) gyration radius of the C$\alpha$ atoms along 100-ns triplicated conventional MD simulation. ConvMD Rep 1 correspond to the trajectory used for analysis in the main text.

### Nucleocapsid at Physiological Concentration(№3) Replicated Conventional Simulations

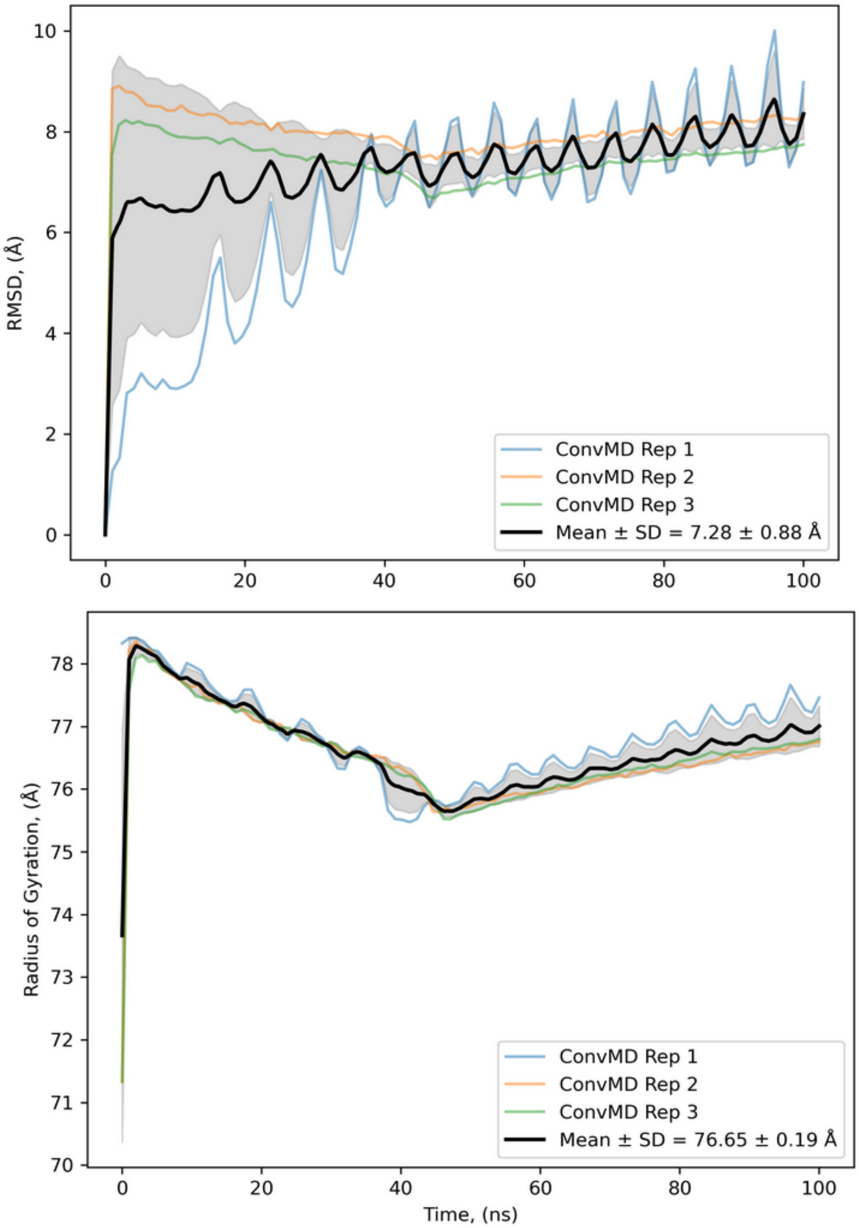


Figure S3. (Top) Root Mean Square Deviation (RMSD) and (bottom) gyration radius of the C$\alpha$ atoms along 100-ns triplicated conventional MD simulation. ConvMD Rep 1 correspond to the trajectory used for analysis in the main text.

### Intermonomeric Salt Bridges

**Table S1. Salt-bridge patterns between consecutive monomers collected at pH=7.**

| Salt Bridge Pair @ pH=7 | Consecutive monomer Pairs |
|---|---|
| ASP15-ARG925 | C0-C14, C15-C29, C20-C19, C30-C44, C35-C34, C40-C39, C45-C59, C5-C4, C50-C49, C55-C54 |
| ASP15-ARG931 | C0-C14, C15-C29, C30-C44, C5-C4, C55-C54 |
| ASP215-ARG325 | C1-C31, C11-C26, C21-C36, C26-C11, C31-C1, C41-C56, C46-C16, C56-C41 |
| ASP215-ARG331 | C1-C31, C21-C36, C41-C56, C46-C16 |
| ASP268-ARG266 | C31-C1, C41-C56 |
| ASP415-ARG731 | C12-C23, C2-C43, C37-C18 |
| ASP415-ARG725 | C22-C33, C47-C28, C52-C3 |
| ASP468-ARG666 | C32-C13, C52-C3 |
| ASP615-ARG525 | C13-C32, C18-C37, C28-C47, C3-C52, C33-C22, C43-C2, C58-C17, C53-C42 |
| ASP615-ARG531 | C13-C32 |
| ASP668-ARG466 | C23-C12, C28-C47, C38-C57 |
| ASP668-ARG439 | C43-C2 |
| ASP68-ARG866 | C25-C24, C35-C34, C40-C39 |
| ASP815-ARG125 | C14-C0, C19-C20, C29-C15, C39-C40, C4-C5, C49-C50, C54-C55 |
| ASP815-ARG131 | C14-C0, C4-C5, C54-C55 |
| ASP868-ARG66 | C29-C15, C49-C50, C54-C55 |
| GLU107-ARG866 | C10-C14, C50-C54 |
| GLU108-ARG866 | C25-C29, C35-C39, C40-C44 |
| GLU110-ARG824 | C15-C29 |
| GLU110-LYS30 | C15-C20, C20-C25, C25-C15, C40-C30 |
| GLU244-ARG512 | C11-C12, C21-C22, C16-C17, C31-C32, C36-C37, C51-C52, C46-C47, C56-C57 |
| GLU290-ARG119 | C1-C0, C26-C25, C56-C55 |
| GLU307-ARG66 | C11-C10, C21-C20, C46-C45 |

| | |
|---|---|
| GLU307-ARG39 | C46-C45 |
| GLU308-ARG66 | C1-C0, C26-C25, C31-C30 |
| GLU310-LYS830 | C31-C44 |
| GLU444-ARG712 | C17-C18, C12-C13, C2-C3, C22-C23, C27-C28, C32-C33, C42-C43, C52-C53, C47-C48, C57-C58 |
| GLU44-ARG312 | C0-C1, C10-C11, C15-C16, C25-C26, C20-C21, C30-C31, C35-C36, C45-C46, C50-C51, C55-C56 |
| GLU490-ARG319 | C17-C16, C42-C41, C57-C56 |
| GLU507-ARG266 | C12-C11, C17-C16, C22-C21, C37-C36, C52-C51 |
| GLU508-ARG266 | C47-C46, C57-C56 |
| GLU644-ARG912 | C18-C19, C13-C14, C23-C24, C3-C4, C43-C44, C38-C39, C53-C54, C58-C59 |
| GLU690-ARG519 | C18-C17, C3-C2, C33-C32, C38-C37 |
| GLU707-ARG466 | C23-C22, C33-C32, C38-C37 |
| GLU710-LYS630 | C18-C58 |
| GLU844-ARG112 | C14-C10, C19-C15, C24-C20, C29-C25, C34-C30, C49-C45, C39-C35 |
| GLU890-ARG719 | C24-C23, C29-C28, C34-C33, C39-C38, C44-C43, C49-C48 |
| GLU907-ARG639 | C34-C33 |
| GLU907-ARG666 | C4-C3 |
| GLU90-ARG919 | C0-C4, C25-C29 |
| GLU910-LYS430 | C29-C47, C34-C22, C44-C2 |

**Table S2. Salt-bridge patterns between consecutive monomers collected at pH=5.**

| Salt Bridge Pair @ pH=5 | Consecutive monomer Pairs |
|---|---|
| GLU107-ARG66 | C0-C2, C8-C9, C7-C8, C6-C7, C5-C6, C4-C0, C59-C55, C58-C59, C57-C58, C56-C57, C55-C56, C54-C50, C53-C54, C51-C52, C52-C53, C50-C51, C49-C45, C48-C49, C3-C4, C47-C48, C46-C47, C45-C46, C44-C40, C43-C44, C42-C43, C41-C42, C40-C41, C39-C35, C2-C3, C38-C39, C37-C38, C36-C37, C35-C36, C33-C34, C34-C30, C32-C33, C31-C32, C29-C25, C30-C31, C28-C29, C1-C2, C26-C27, C27-C28, C25-C26, C24-C20, C23-C24, C22-C23, C21-C22, C20-C21, C19-C15, C18-C19, C17-C18, C16-C17, C14-C10, C15-C16, C13-C14, C12-C13, C10-C11, C11-C12, C9-C5 |
| GLU44-ARG112 | C8-C7, C7-C6, C6-C5, C5-C9, C59-C58, C4-C3, C58-C57, C57-C56, C56-C55, C55-C59, C54-C53, C53-C52, C52-C51, C51-C50, C50-C54, C49-C48, C3-C2, C48-C47, C47-C46, C46-C45, C45-C49, C44-C43, C43-C42, C42-C41, C41-C40, C40-C44, C39-C38, C2-C1, C38-C37, C37-C36, C36-C35, C35-C39, C34-C33, C33-C32, C32-C31, C31-C30, C30-C34, C29-C28, C1-C0, C28-C27, C27-C26, C26-C25, C25-C29, C24-C23, C23-C22, C22-C21, C21-C20, C20-C24, C19-C18, C18-C17, C17-C16, C16-C15, C15-C19, C14-C13, C13-C12, C12-C11, C11-C10, C10-C14, C9-C8, C0-C5 |
| ASP15-ARG125 | C0-C6, C8-C54, C7-C56, C6-C32, C5-C0, C59-C18, C4-C33, C57-C31, C58-C39, C56-C7, C55-C50, C54-C8, C53-C24, C51-C17, C52-C26, C50-C55, C49-C3, C48-C34, C3-C49, C47-C36, C46-C12, C44-C13, C45-C40, C43-C29, C42-C21, C41-C2, C40-C45, C39-C58, C2-C41, C38-C19, C37-C11, C36-C47, C35-C30, C34-C48, C33-C4, C31-C57, C32-C6, C30-C35, C29-C43, C28-C14, C1-C22, C27-C16, C26-C52, C25-C20, C24-C53, C23-C9, C22-C1, C20-C25, C21-C42, C19-C38, C18-C59, C16-C27, C17-C51, C15-C10, C14-C28, C13-C44, C12-C46, C11-C37, C10-C15, C9-C23 |

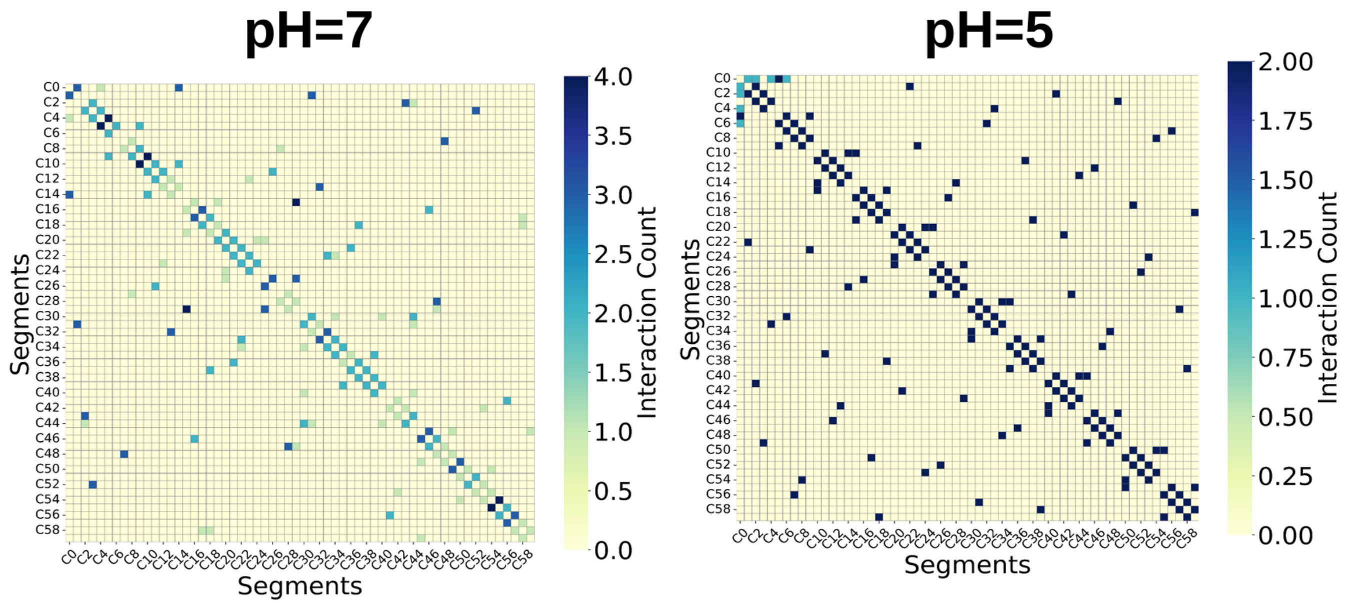


Figure S4. Intermonomeric salt bridge interaction pattern map for STMV particles at pH=7 (№1-5) and at pH=5 (№6).

### Precisions on Gyration Radius from Conventional MD

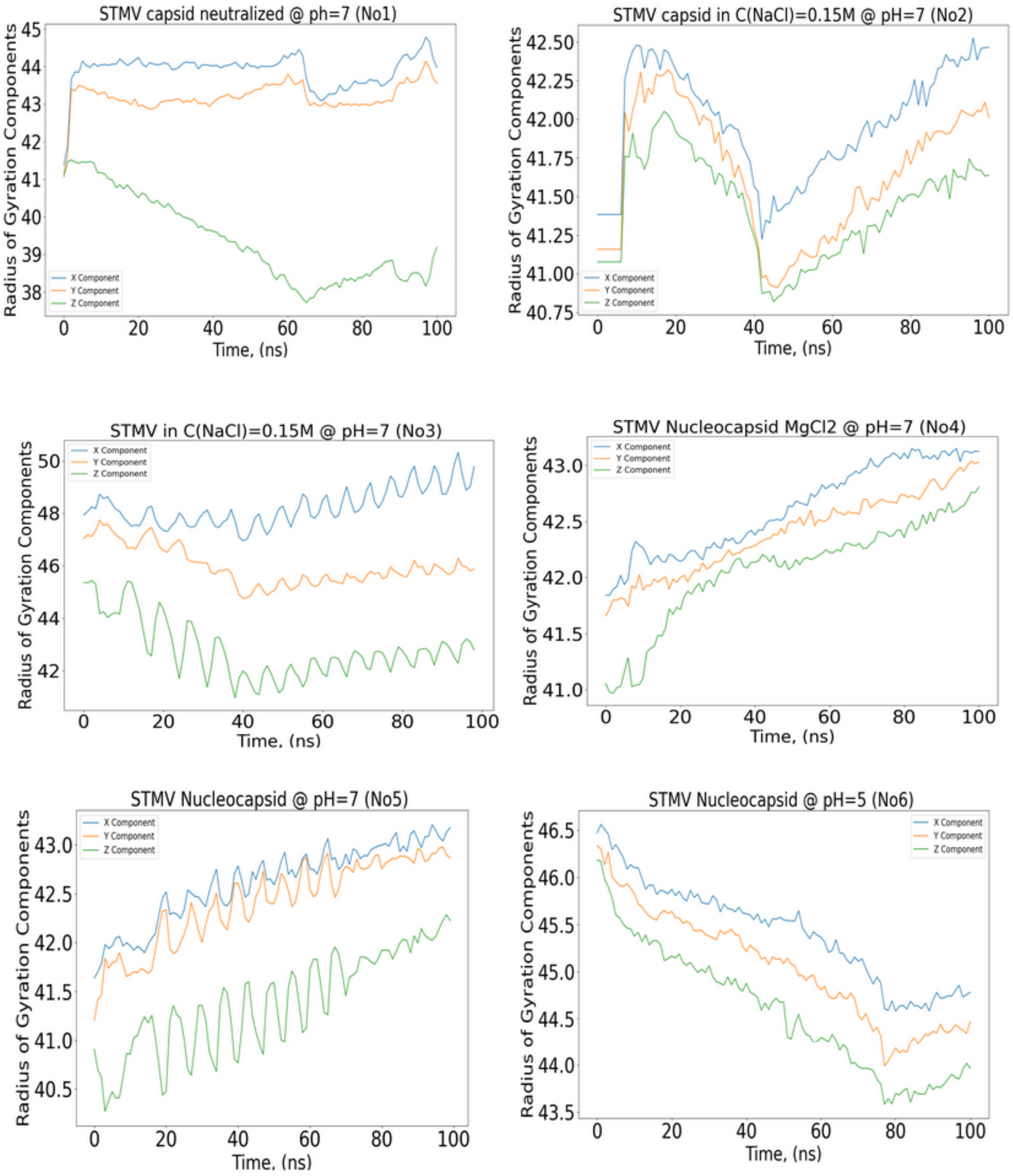


Figure S5. The principal radii of gyration $r_{gyr,x}$, $r_{gyr,y}$, and $r_{gyr,z}$ in blue, orange, and green.

### Details on RMSF of Conventional MD

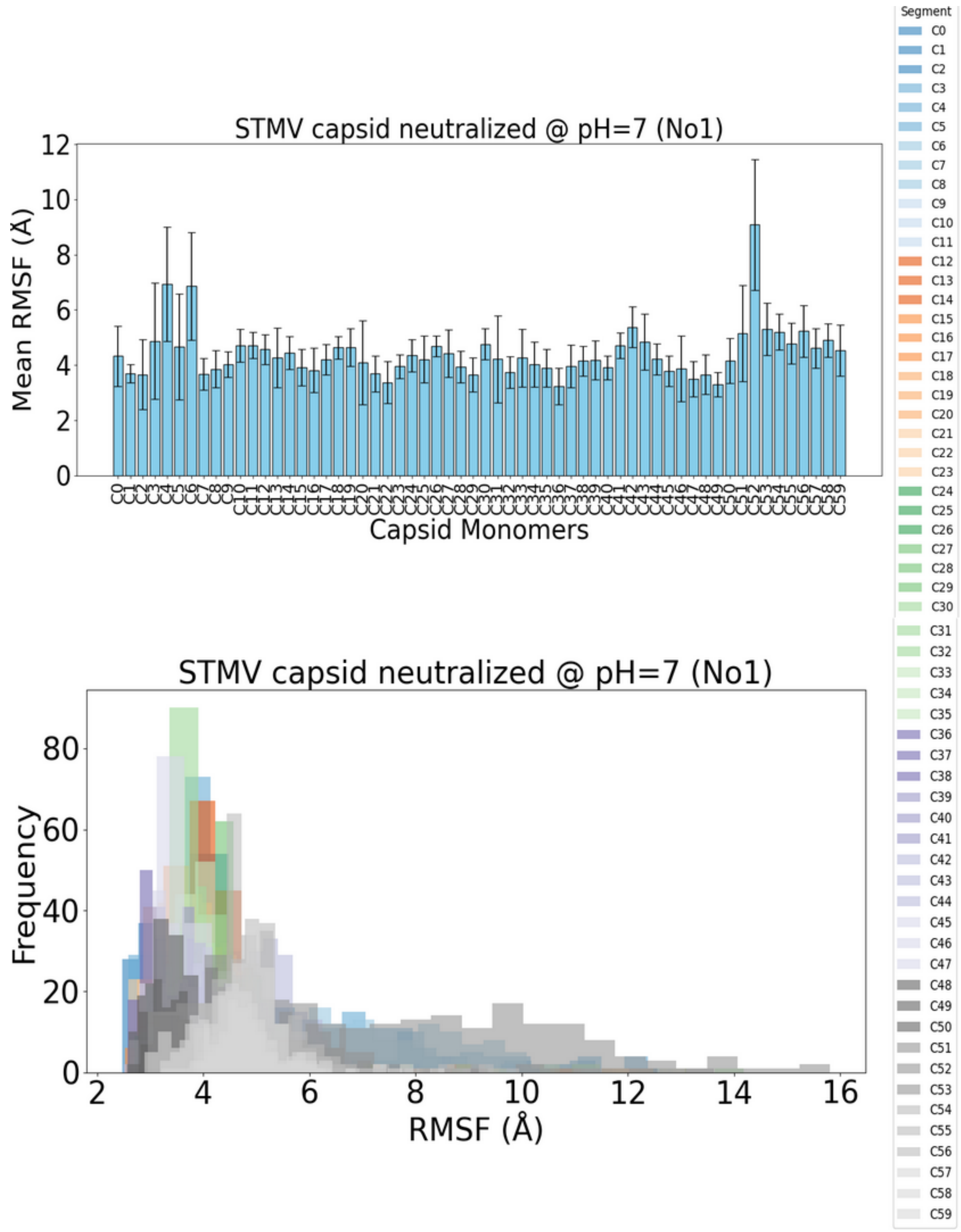

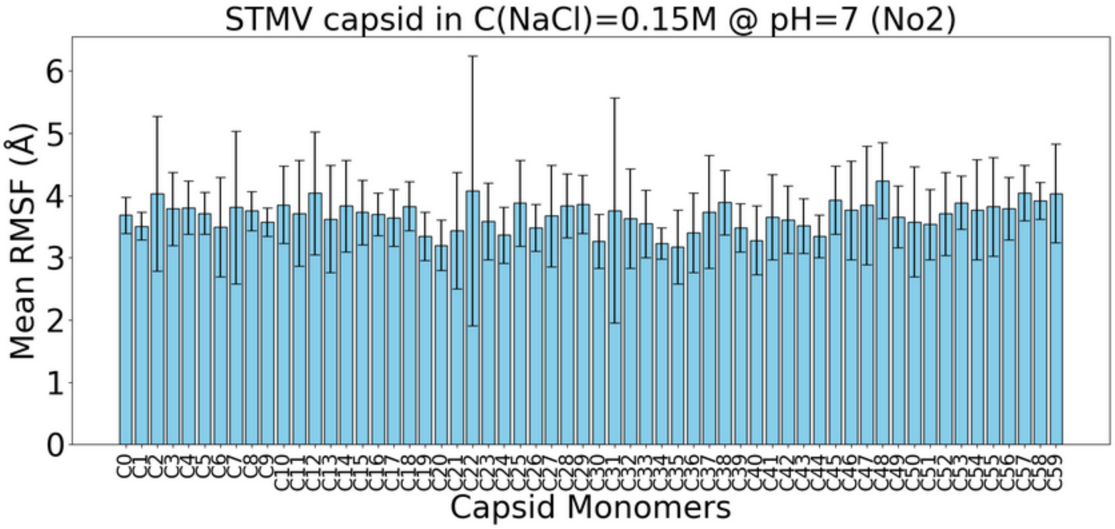
STMV capsid in C(NaCl)=0.15M @ pH=7 (No2)
Mean RMSF (Å)
0
1
2
3
4
5
6
C0
C1
C2
C3
C4
C5
C6
C7
C8
C9
C10
C11
C12
C13
C14
C15
C16
C17
C18
C19
C20
C21
C22
C23
C24
C25
C26
C27
C28
C29
C30
C31
C32
C33
C34
C35
C36
C37
C38
C39
C40
C41
C42
C43
C44
C45
C46
C47
C48
C49
C50
C51
C52
C53
C54
C55
C56
C57
C58
C59
Capsid Monomers

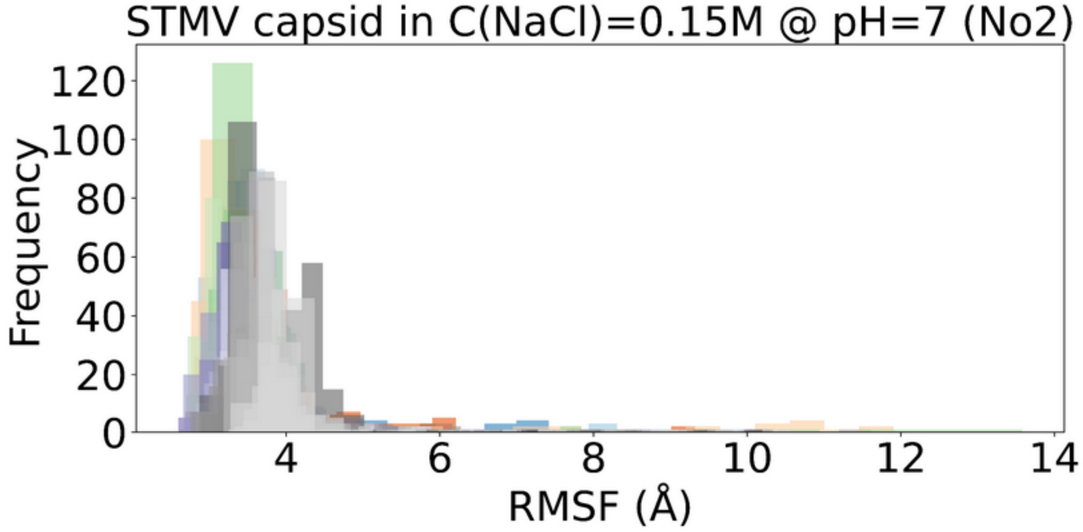
STMV capsid in C(NaCl)=0.15M @ pH=7 (No2)
Frequency
0
20
40
60
80
100
120
4
6
8
10
12
14
RMSF (Å)

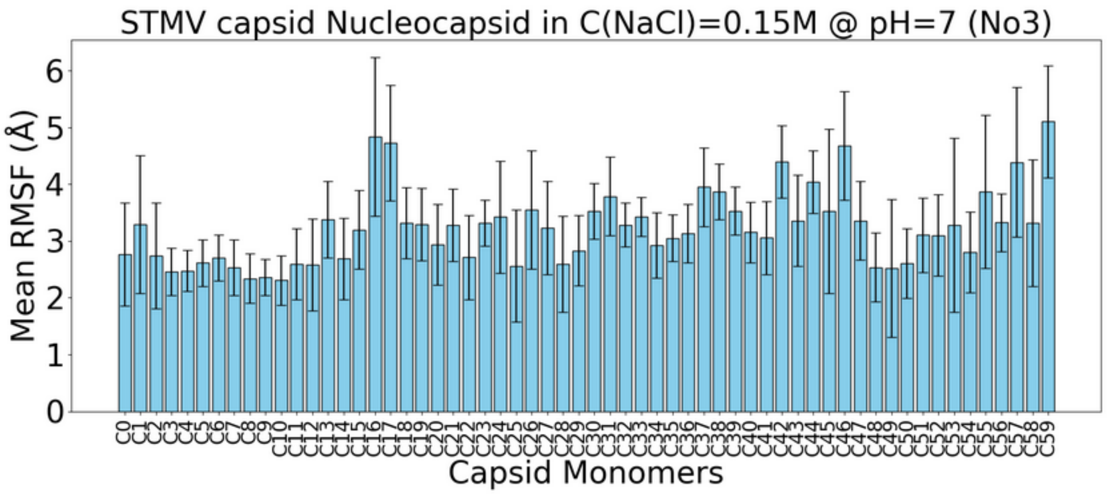
STMV capsid Nucleocapsid in C(NaCl)=0.15M @ pH=7 (No3)
Mean RMSF (Å)
Capsid Monomers
C0 C1 C2 C3 C4 C5 C6 C7 C8 C9 C10 C11 C12 C13 C14 C15 C16 C17 C18 C19 C20 C21 C22 C23 C24 C25 C26 C27 C28 C29 C30 C31 C32 C33 C34 C35 C36 C37 C38 C39 C40 C41 C42 C43 C44 C45 C46 C47 C48 C49 C50 C51 C52 C53 C54 C55 C56 C57 C58 C59
0 1 2 3 4 5 6

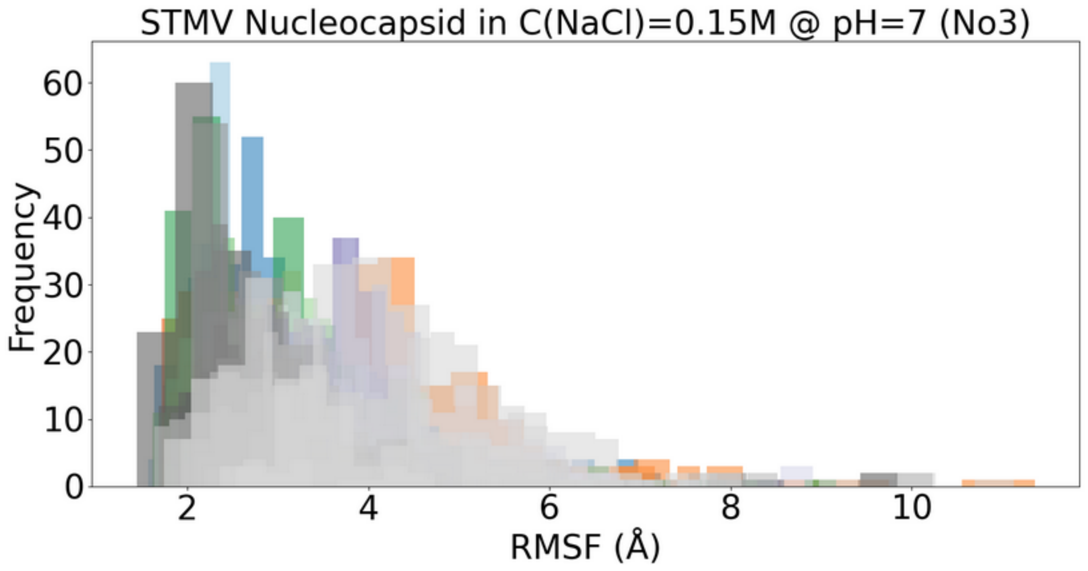
STMV Nucleocapsid in C(NaCl)=0.15M @ pH=7 (No3)
Frequency
RMSF (Å)
0 10 20 30 40 50 60
2 4 6 8 10

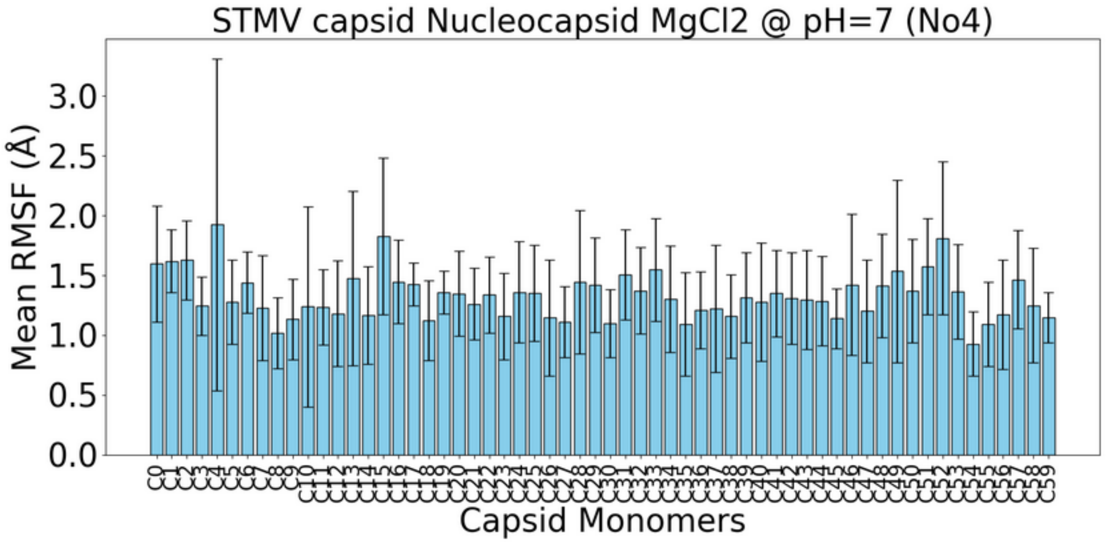
STMV capsid Nucleocapsid MgCl2 @ pH=7 (No4)
Mean RMSF (Å)
Capsid Monomers

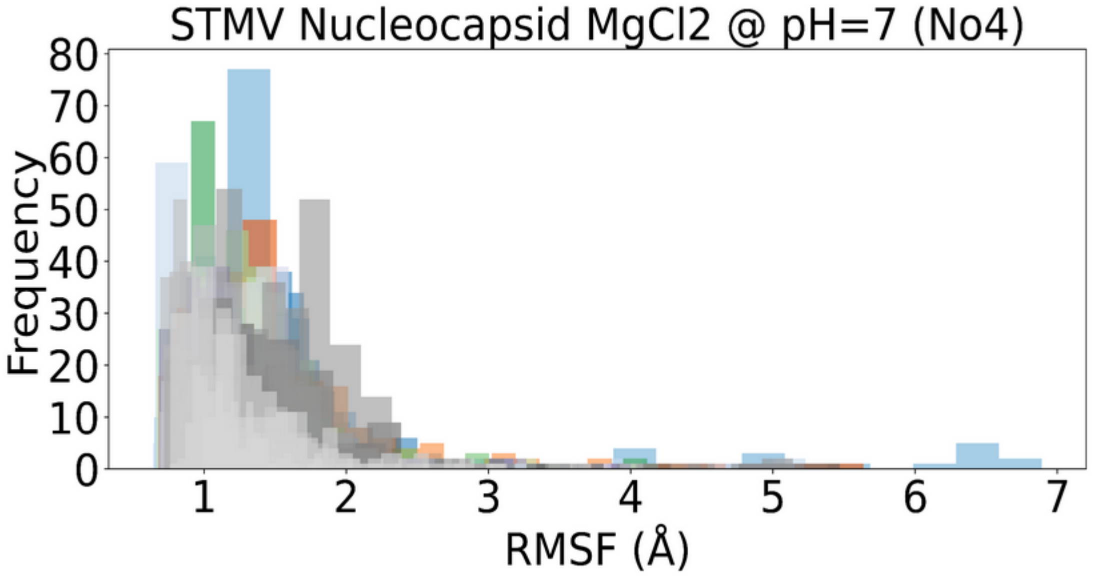
STMV Nucleocapsid MgCl2 @ pH=7 (No4)
Frequency
RMSF (Å)

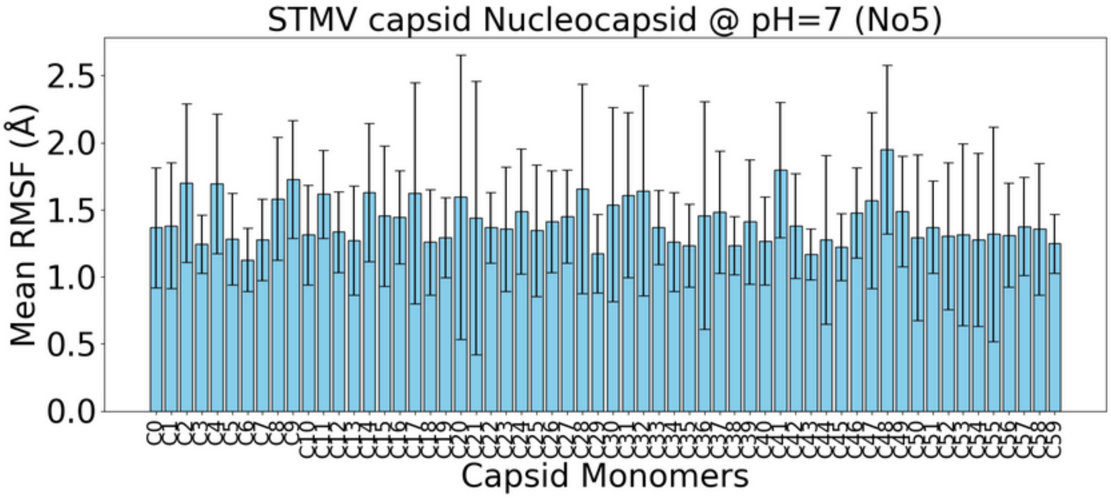
STMV capsid Nucleocapsid @ pH=7 (No5)
Mean RMSF (Å)
Capsid Monomers

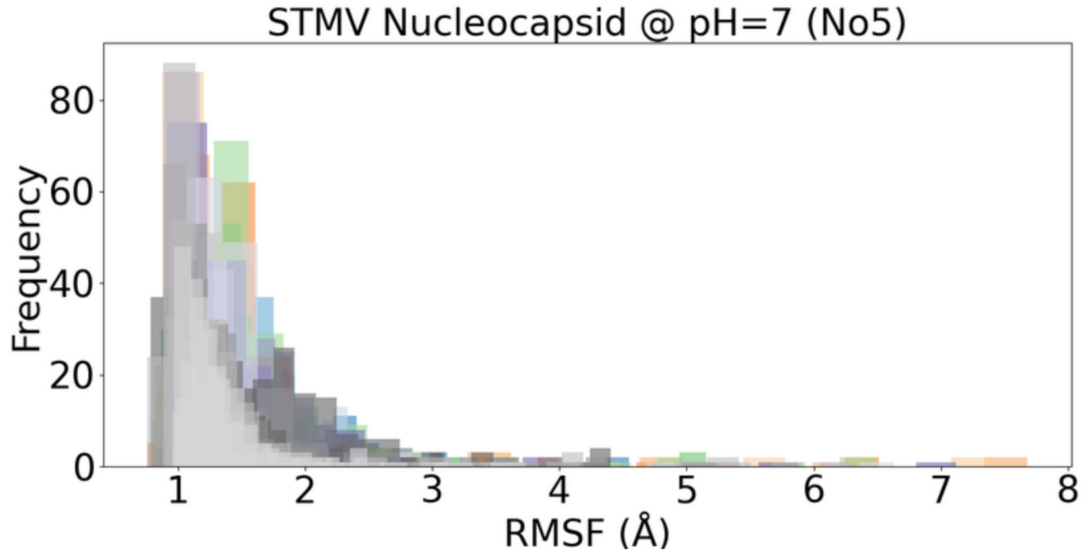
STMV Nucleocapsid @ pH=7 (No5)
Frequency
RMSF (Å)

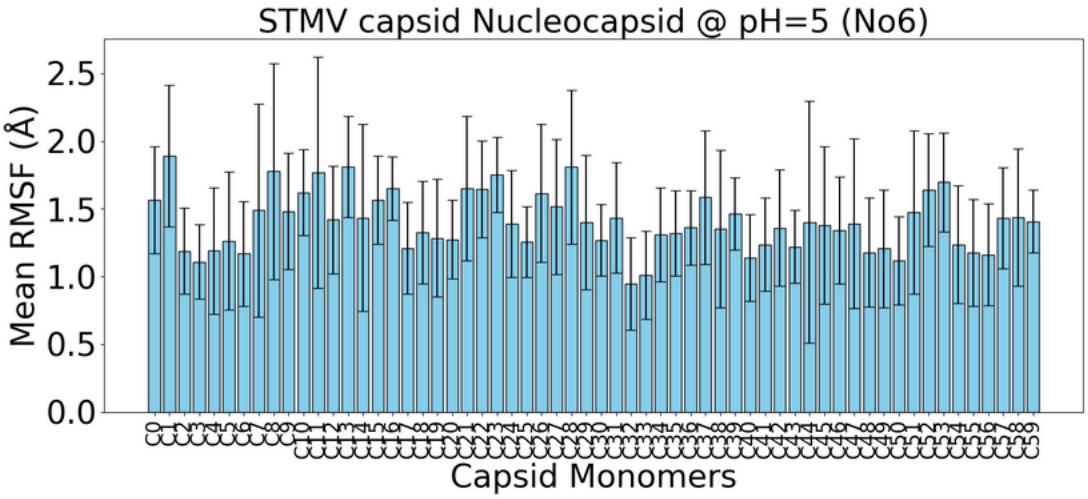


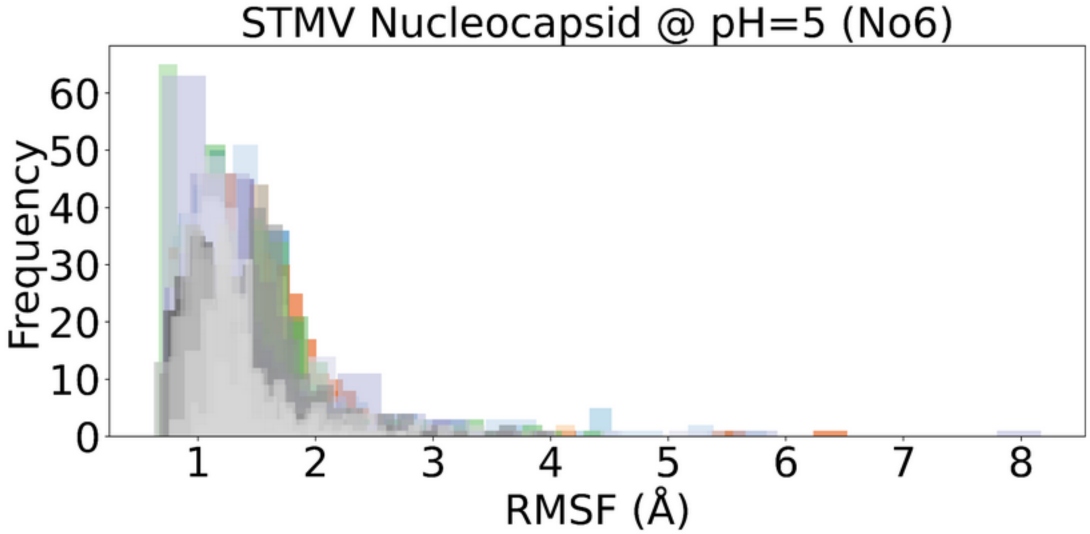


Figure S6. (Top) Monomeric RMSF with computed mean and standard deviation for each capsomer. (Bottom) RMSF distribution for each monomer collected for each assembly.

### Capsid-RNA Hydrogen Bond Interactions

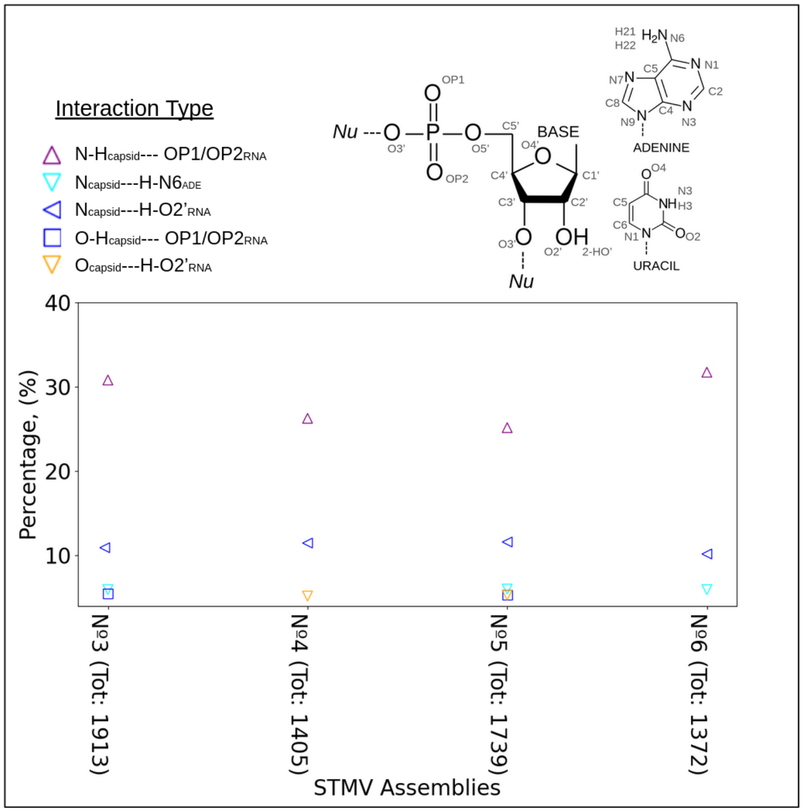


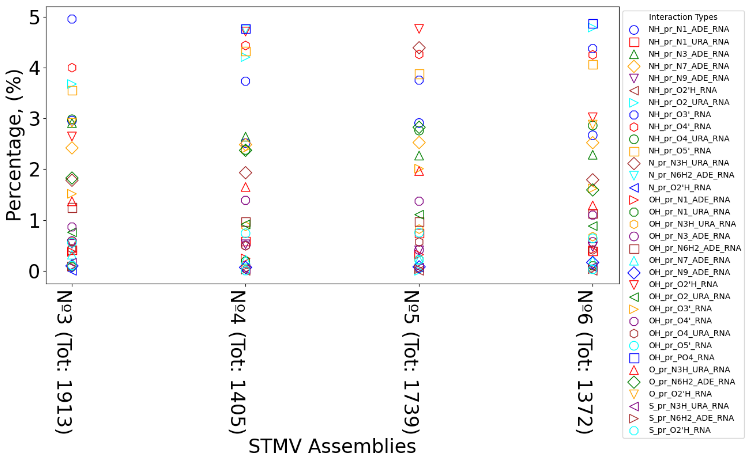


Figure S7. Percentage contribution of interaction types accounting for more than 5% (top) and less than 5 % (bottom) of total hydrogen bonds in each system throughout the simulation, focusing on hydrogen bonds between the capsid protein core and nucleotide groups across four STMV nucleocapsid-related particles depicted here as STMV assemblies. 'Tot' represents the total hydrogen bonds observed in the MD simulations, with 'Nu' referring to the nucleotide. Donors are marked with '-H', and ADE and URA correspond to adenine and uracil bases of the ssRNA.

### Capsid-RNA Salt Bridges

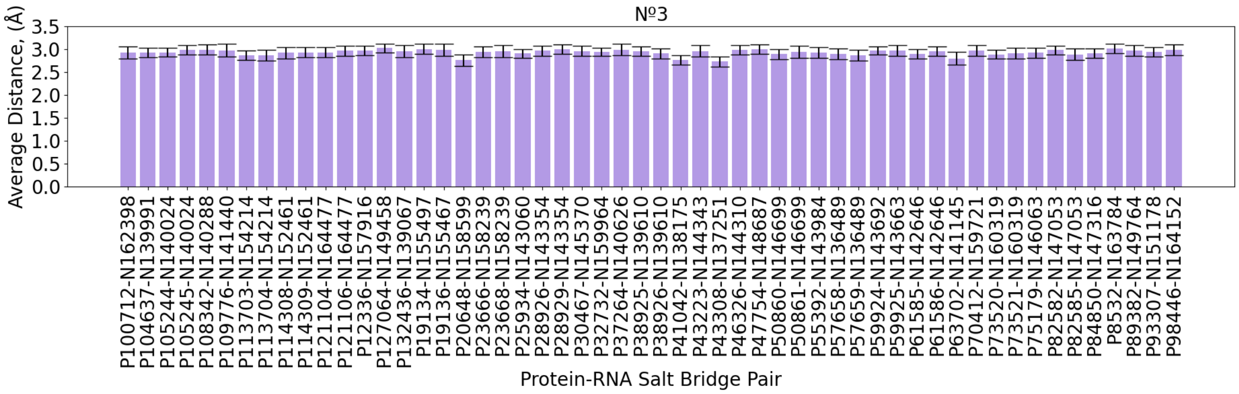

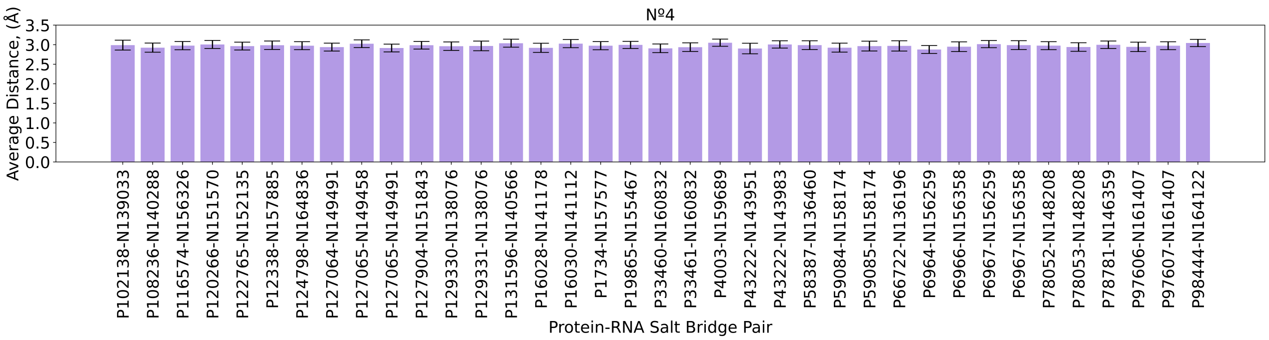
№4
Average Distance, (Å)
3.5
3.0
2.5
2.0
1.5
1.0
0.5
0.0
P102138-N139033
P108236-N140288
P116574-N156326
P120266-N151570
P122765-N152135
P12338-N157885
P124798-N164836
P127064-N149491
P127065-N149458
P127065-N149491
P127904-N151843
P129330-N138076
P129331-N138076
P131596-N140566
P16028-N141178
P16030-N141112
P1734-N157577
P19865-N155467
P33460-N160832
P33461-N160832
P4003-N159689
P43222-N143951
P43222-N143983
P58387-N136460
P59084-N158174
P59085-N158174
P66722-N136196
P6964-N156259
P6966-N156358
P6967-N156259
P6967-N156358
P78052-N148208
P78053-N148208
P78781-N146359
P97606-N161407
P97607-N161407
P98444-N164122
Protein-RNA Salt Bridge Pair

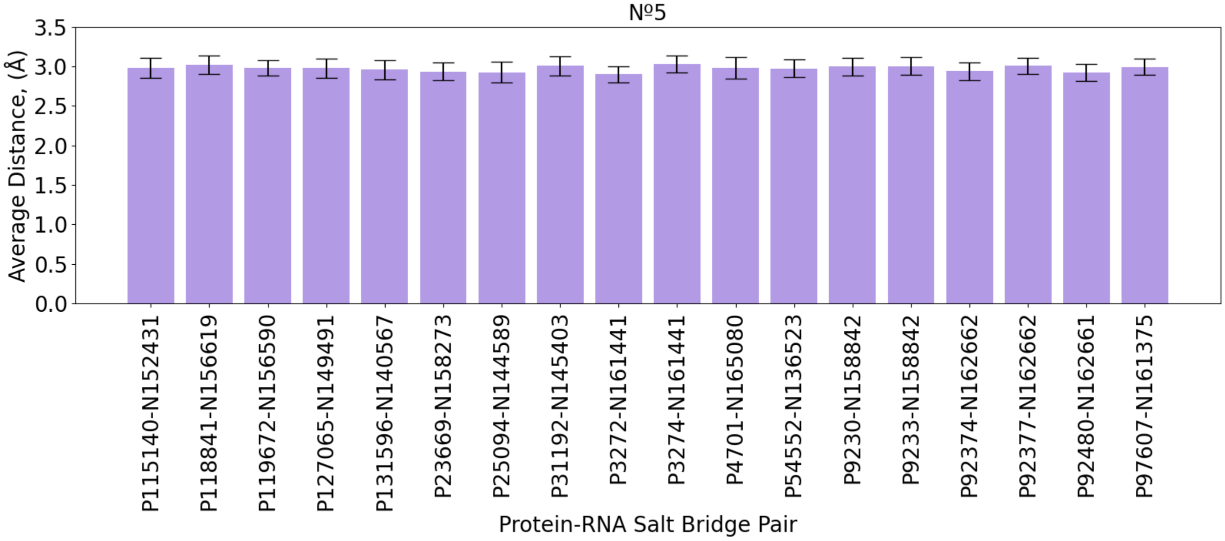

№5
Average Distance, (Å)
3.5
3.0
2.5
2.0
1.5
1.0
0.5
0.0
P115140-N152431
P118841-N156619
P119672-N156590
P127065-N149491
P131596-N140567
P23669-N158273
P25094-N144589
P31192-N145403
P3272-N161441
P3274-N161441
P4701-N165080
P54552-N136523
P9230-N158842
P9233-N158842
P92374-N162662
P92377-N162662
P92480-N162661
P97607-N161375
Protein-RNA Salt Bridge Pair

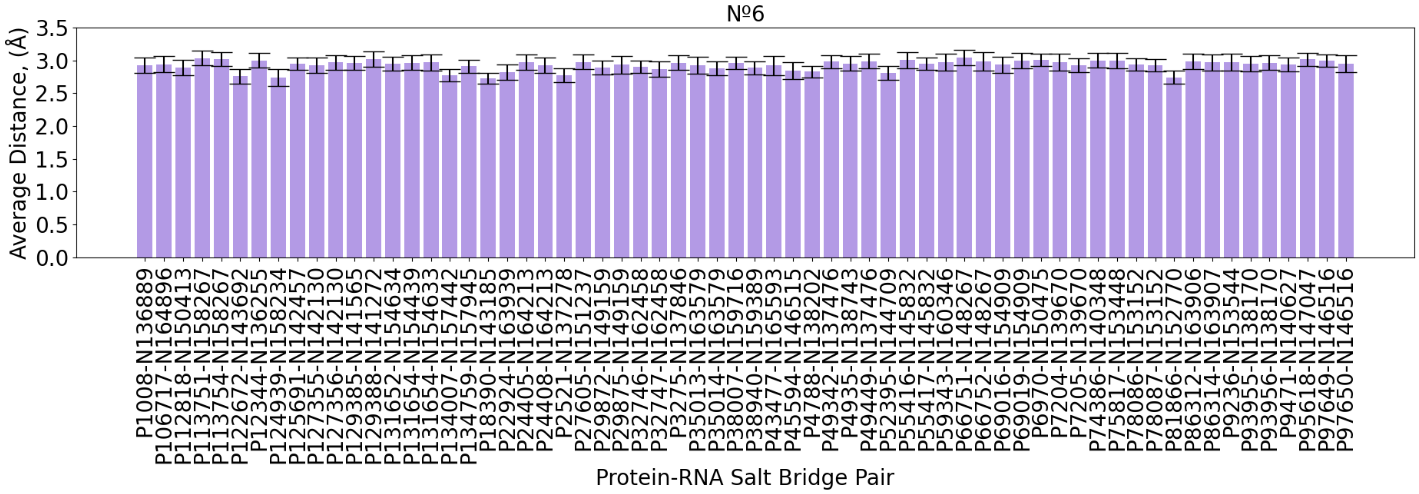


Figure S8. Stable salt bridge interactions (present more than 30 % of the simulation) between positively charged residues of capsid (ARG and LYS charged ammonium atoms) and oxygens of phosphate groups of RNA.

### Dual-Boost Gaussian Accelerated Dynamics

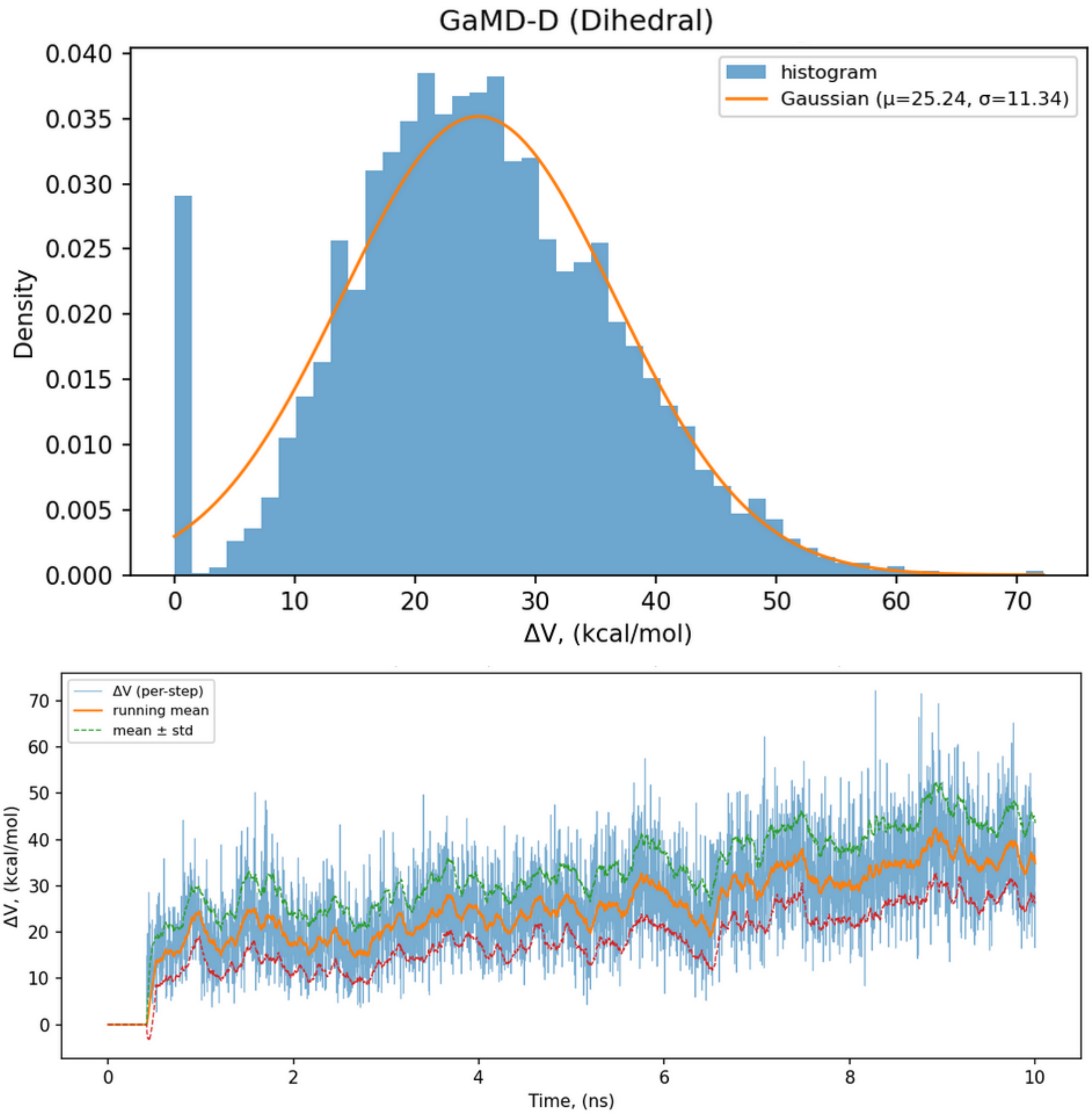


Figure S9. (Top) Distribution of the dihedral boost potential ($\Delta$V) during the GaMD simulation. The histogram (blue) and fitted Gaussian (orange) confirm that the boost potential follows an approximately normal distribution ($\mu$ = 25.0 kcal/mol, $\sigma$ = 11.1 kcal/mol). (Bottom) Time evolution of $\Delta$V along the GaMD simulation. The instantaneous $\Delta$V (blue) fluctuates around a stable mean (orange line) with consistent variance (dashed lines, $\pm\sigma$), indicating equilibrium and proper boost potential behavior.

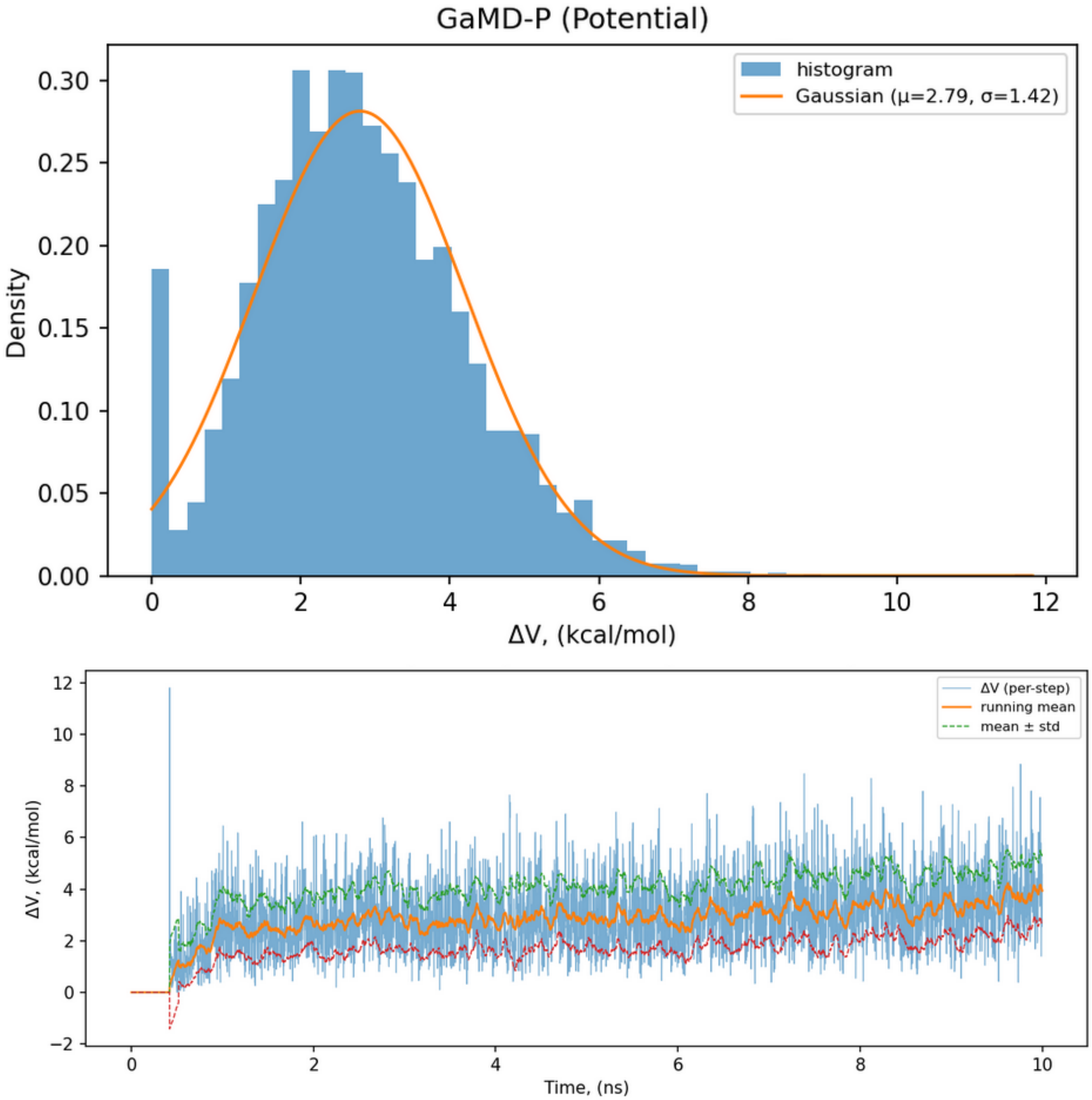


Figure S10. (Top) Distribution of the potential boost potential ($\Delta$V) during the GaMD simulation. The nearly Gaussian shape ($\mu$ = 2.7 kcal/mol, $\sigma$ = 0.6 kcal/mol) and narrow width confirm that the applied boost meets the target standard deviation ($\sigma$ = 3 kcal/mol). (Bottom) Time evolution of $\Delta$V along the GaMD simulation. The boost potential remains statistically stationary, demonstrating a well-behaved and stable boost during production.

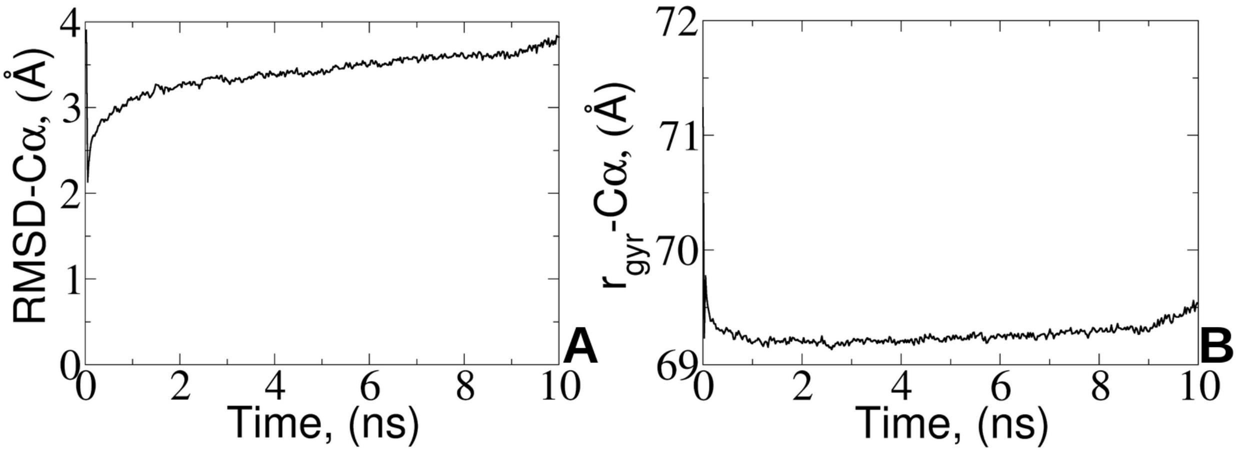


Figure S11. Time evolution of C$\alpha$ RMSD (left) and radius of gyration (right) of the STMV capsid from a 10-ns dual-boost GaMD simulation of assembly № 2, corresponding to an effective sampling of up to microsecond-equivalent of conventional MD.

### Example of Colvars File for PMF Calculation within WTM-eABF Approach

```
colvarsTrajFrequency      5000
colvarsRestartFrequency   5000
indexFile                 backbone_carbon_serials.ndx

colvar {
    name R_gyr

    width 1

    lowerboundary  65.0
    upperboundary  85.0

gyration {
        atoms {
            indexGroup  capsid_carbons
        }
    }
}


metadynamics {
    colvars                R_gyr
    hillWidth              5.0
    hillWeight             0.1
    wellTempered           on
    biasTemperature        3000
    writeFreeEnergyFile on
    multipleReplicas on
    replicasRegistry replica_registry.txt
```

```
    replicaUpdateFrequency 5000
    writePartialFreeEnergyFile on
    keepFreeEnergyFiles on

}
```

### PMF Histograms

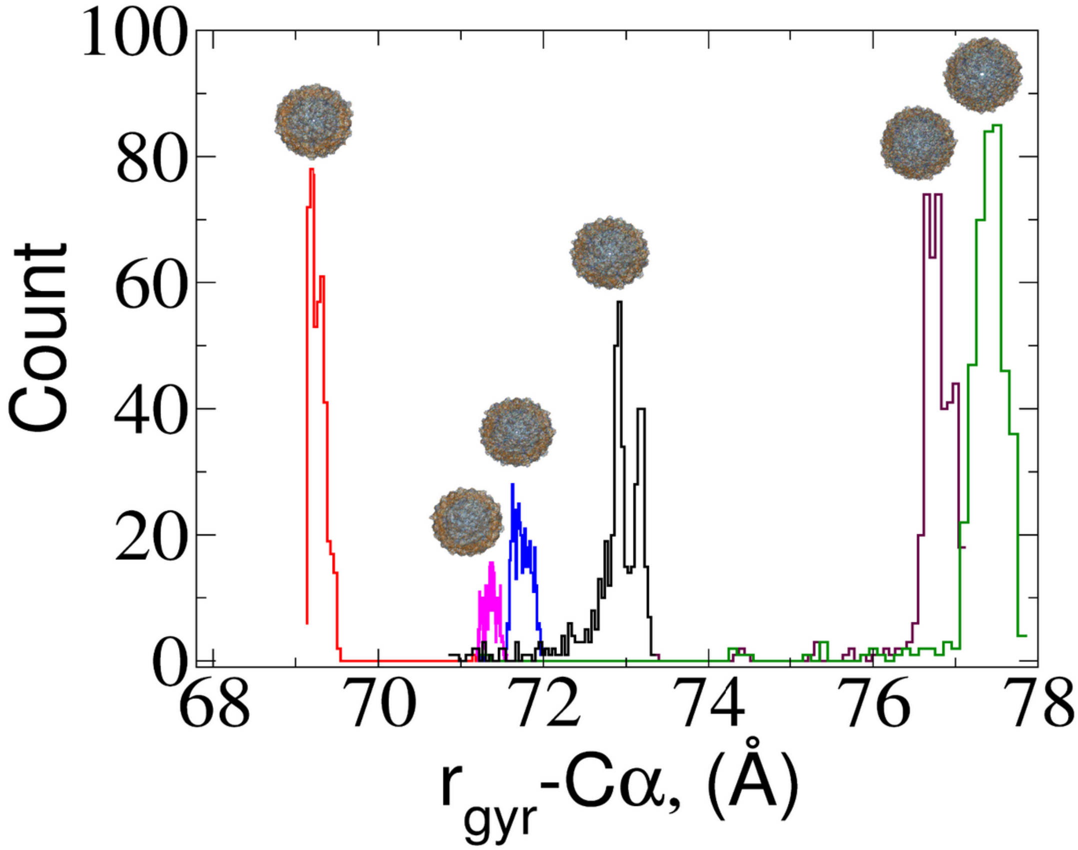


Figure S12. Histograms of the related WTMetaD PMF calculations along the radius of gyration $r_{gyr}$ for the STMV capsid: №1 - neutralized capsid with $Cl^-$ ions (black), №2 - capsid at physiological NaCl concentration (red), №3 - nucleocapsid at physiological NaCl concentration (green), №4 - nucleocapsid neutralized with $MgCl_2$ (blue), №5 - nucleocapsid at physiological NaCl concentration with $Mg^{2+}$ present (magenta), and №6 - nucleocapsid in 0.5 M NaCl concentration with $Mg^{2+}$ mimicking pH=5 conditions (brown). The structure of each capsid with the gradient directions for each $C_\alpha$ atom corresponding to the highest peaks of the histograms are depicted for each particle.

#### Capsid Accessibility and Porosity

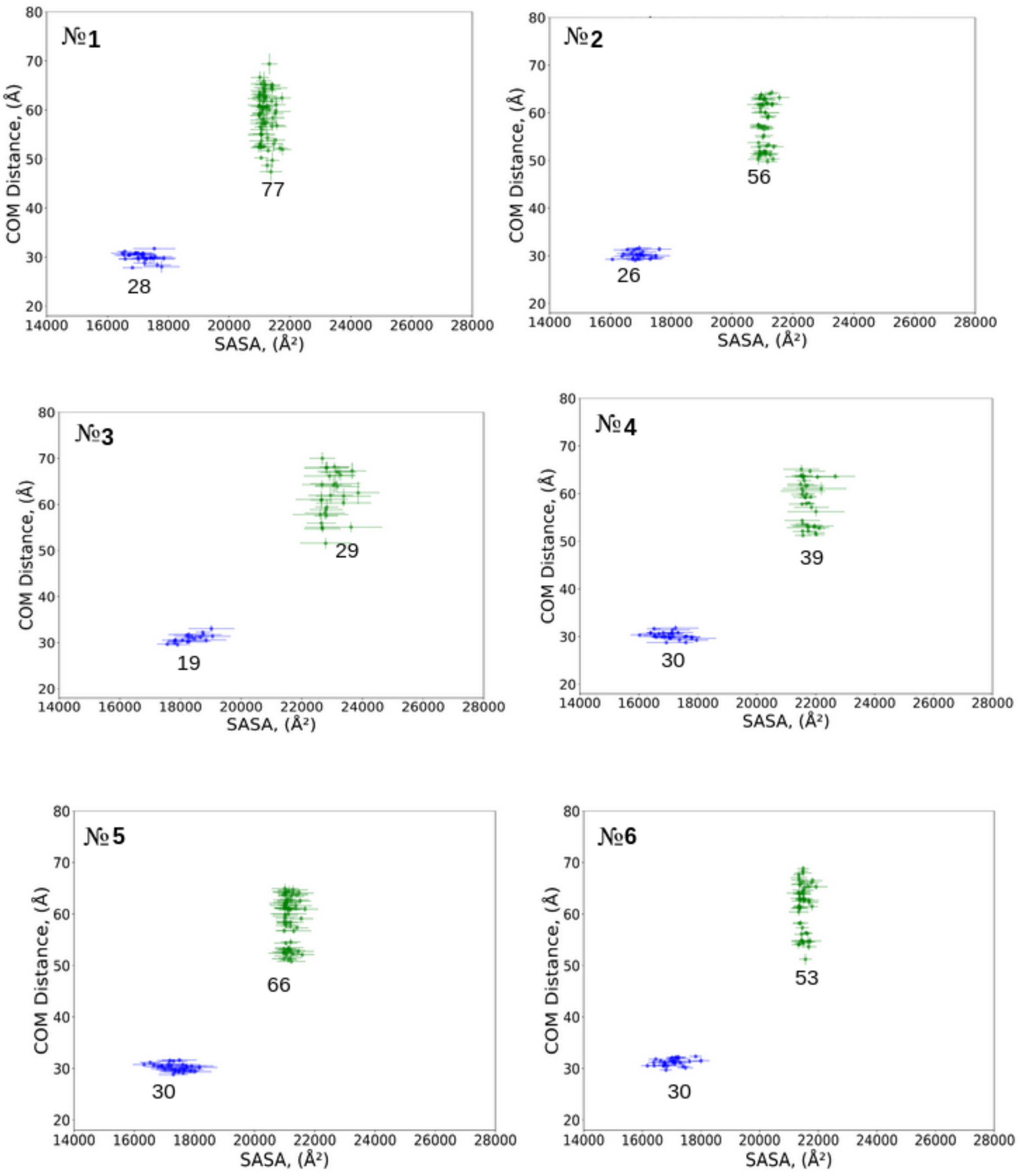


Figure S13. K-means clustering along two variables, COM distance between consecutive monomers and SASA of these monomeric pairs. The occluded pairs are shown in blue, and the pairs creating the intermonomeric pores are shown in green. The total number of pairs and the level of deviation are assigned for each cluster.

#### STMV Permeability

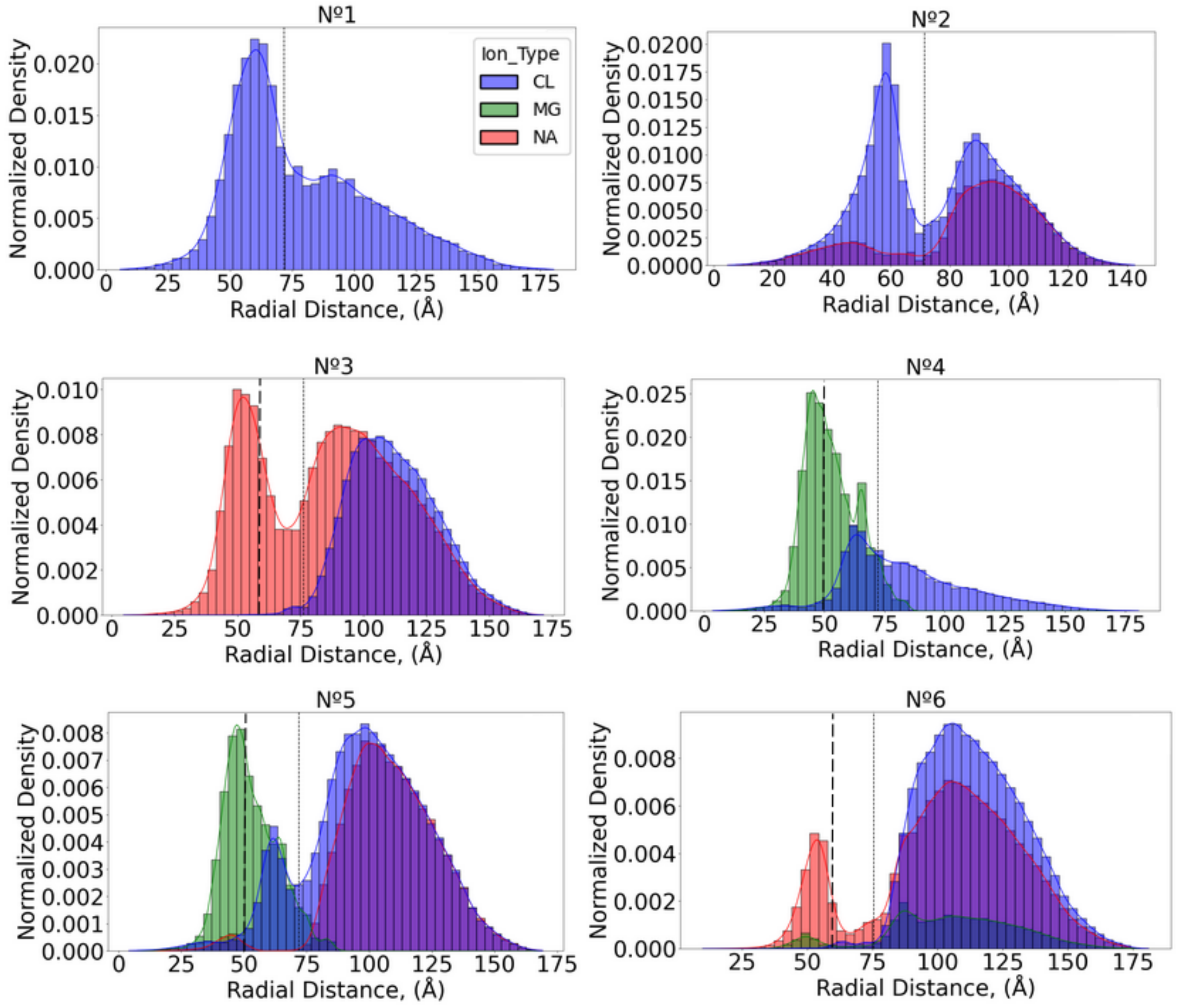


Figure S14. Radial density distribution of ions along 100-ns conventional MD simulation. The thin and bold dashed lines correspond to the average capsid and RNA radii, respectively, illustrated for visual support of the ions localization. The ions $Cl^{-}$, $Mg^{2+}$, and $Na^{+}$ are color-coded in blue, green, and red, respectively.

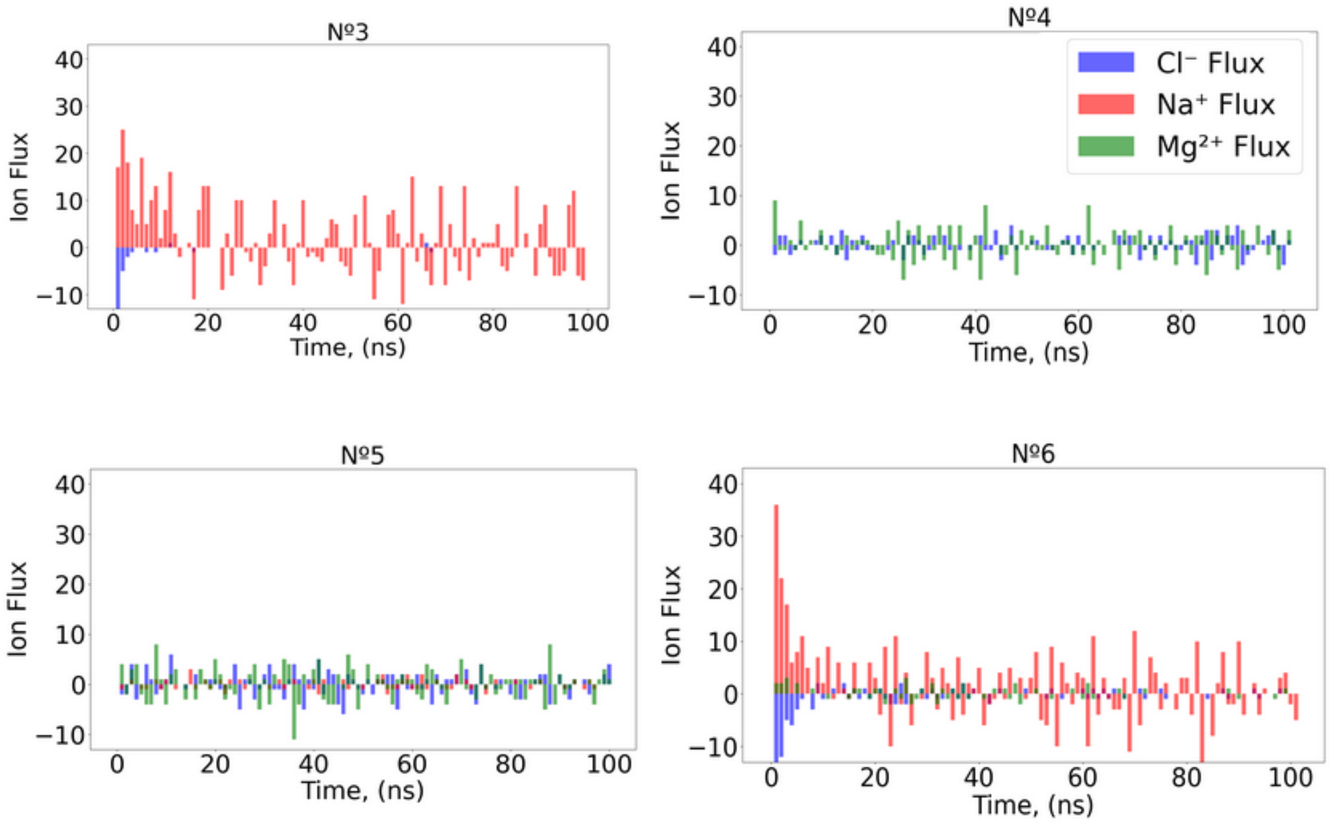


Figure S15. Net Ion flux of $Cl^{-}$, $Na^{+}$, and $Mg^{2+}$ displayed in blue, red, and green, respectively for each of the RNA-containing STMV assemblies of №3–6.

#### RDFs Between Ions and Charged Groups

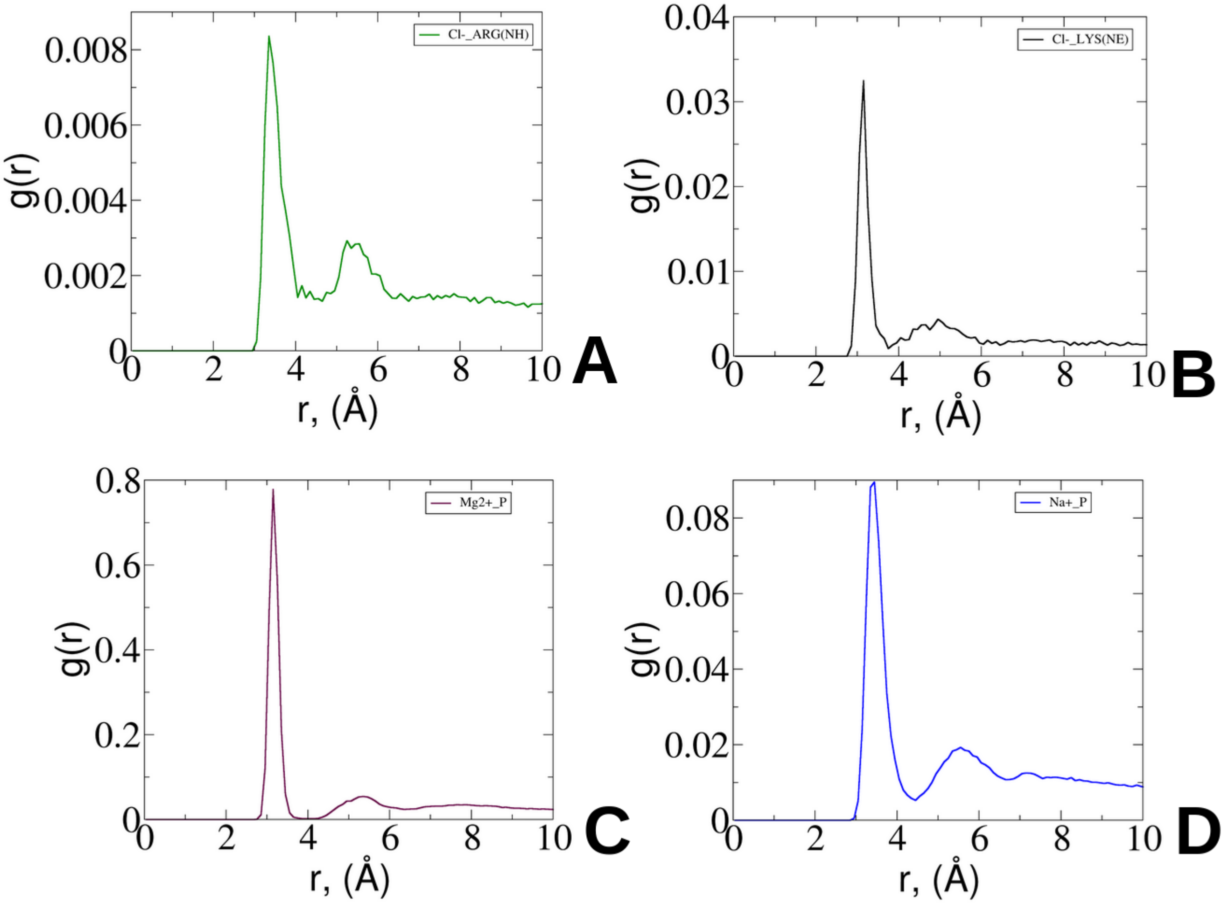


Figure S16. Non-normalized radial distribution functions (RDFs) for $Cl^{-}$ ions interacting with (A) arginine (ARG) side-chain, (B) side-chain lysine (LYS) charged groups. Phosphorus atom of phosphate group of RNA interactions with (C) $Mg^{2+}$ and (D) with $Na^{+}$.

**Residence Time**

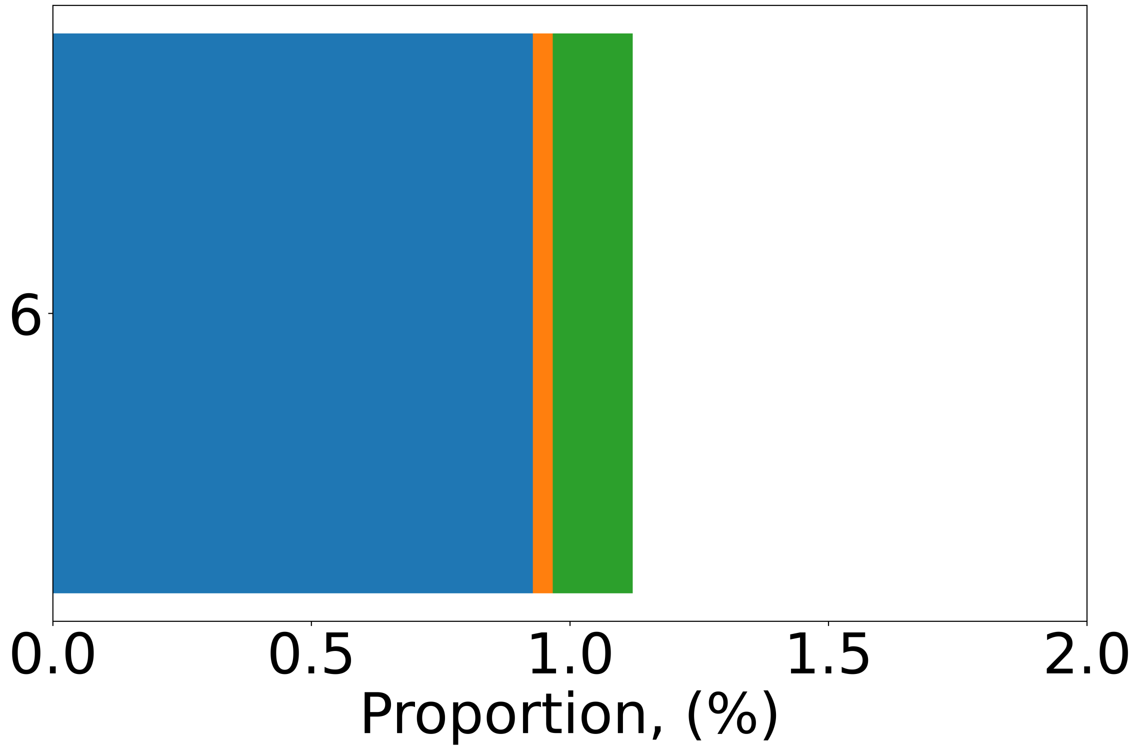


Figure S17. Residence time of $Na^+$ ions interacting with negatively charged side-chains of glutamic acid observed for №6 at pH=5.

**Residual Dipole Moments**

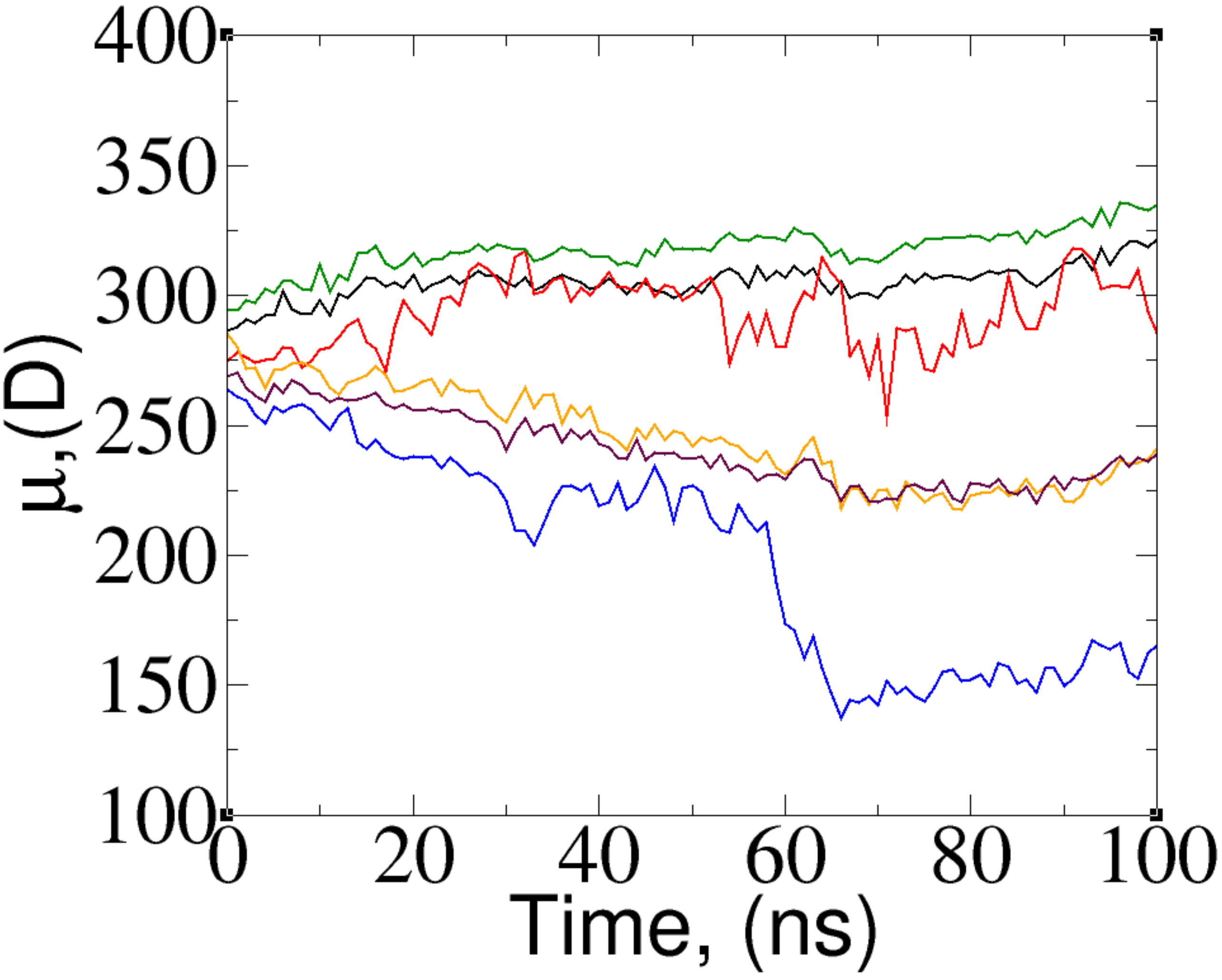


Figure S18. Residual dipole moments evolution along the trajectory upon sal-bridge destruction: $ARG^{C5}_{125}$ - black, $ASP^{C4}_{815}$ - red, $ARG^{C5}_{131}$ - green, $ASP^{C5}_{15}$ - blue, $ARG^{C4}_{931}$ - orange, $ARG^{C4}_{925}$ - brown.
